# Multimodal assessment of acute stress dynamics using an Aversive Video Paradigm (AVP)

**DOI:** 10.1101/2024.04.05.588254

**Authors:** Sumit Roy, Yan Fan, Mohsen Mosayebi-Samani, Maren Claus, Nilay Mutlu, Thomas Kleinsorge, Michael A. Nitsche

**Affiliations:** Department of Psychology and Neurosciences, Leibniz Research Centre for Working Environment and Human Factors, Dortmund, Germany; International Graduate School of Neuroscience (IGSN), Ruhr University Bochum, Bochum, Germany; Department of Immunology, Leibniz Research Centre for Working Environment and Human Factors, Dortmund, Germany; Bioengineering Department, Yildiz Technical University, Istanbul, Turkey; German Centre for Mental Health; Bielefeld University, University Hospital OWL, Protestant Hospital of Bethel Foundation, University Clinic of Psychiatry and Psychotherapy and University Clinic of Child and Adolescent Psychiatry and Psychotherapy

## Abstract

This study explored the efficacy of inducing stress through aversive video clips and investigated its impact on psychological processes, brain, and vegetative physiology. It had a randomized, single-blinded, crossover design, where participants were exposed in separate sessions to aversive or neutral video clips. Subjective feelings of stress were assessed via questionnaires. Electroencephalography (EEG) with 62 electrodes was recorded continuously. EEG power and connectivity changes based on coherence were analyzed. Heart rate (HR) and heart rate variability (HRV) data were obtained during the whole experiment, and saliva was collected for cortisol and cytokine analysis at different time intervals. Subjective data showed increased anxiety and negative affect induced by the aversive video clips, accompanied by elevated salivary cortisol levels after exposure to the stressful clips, and decreased heart rate variability. Cytokine levels however increased over time in both control and stress conditions, which argues against a stress-specific alteration of cytokines in this specific stress protocol. EEG alterations during stress induction suggest a disruption of top-down control and increased bottom-up processing. These results show that aversive video clips are suited to induce psychological stress in an experimental setting reliably, and are associated with stress-specific emotional, and physiological changes.

## Introduction

The issue of stress has gained increasing prominence in modern society, as it exerts profound effects on both, physical and mental well-being, and consequently, on society ^1^. These effects extend beyond the realm of cognition, and emotion, extending their implications to physiology and metabolism of the human body^2^. Psychological stress can be defined as a state where the individual is challenged by a potential loss of control due to external and internal triggers and experiences an increase in allostatic load^3^. Psychological stressors enhance the activity of the hypothalamic-pituitary-adrenal (HPA) axis and the sympathetic nervous system (SNS) ^4^. Increased activity of the SNS and HPA axis leads to physiological changes in multiple organs, including the cardiac system ^5^, endocrine secretion ^6^, brain oscillations and activation ^7,8^, the immune system ^9^, and cognition and behavior ^10^. Thus, stress exerts its impact on diverse systems and warrants a detailed study of these systems.

Stress is a highly dynamic process, particularly concerning acute stressors ^7^. Hence, investigating stress dynamics in multiple modalities promises to provide a comprehensive understanding of how various systems interact synergistically to produce stress effects.

To study stress in a controlled laboratory setting, reliable methods for stress induction are essential. Numerous established methods, such as the Trier Social Stressor task (TSST) and the Montreal Imaging Stressor Task (MIST), have been proven to reliably induce stress, typically involving social evaluative threats ^11,12^. However, these paradigms often demand extensive training, require a relevant amount of research staff members, and it is not trivial in each case to introduce adequate control conditions. To mimic constant stress exposure with uncontrollable and aversive stimuli, which are relevant in modern society, we sought to focus on more selective *psychological* stress induction by showing aversive movie clips. In this paradigm, participants are exposed to threatening stimuli through aversive videos, without the need for active engagement or social evaluation ^13,14^. Such stress paradigms align with the contemporary stress landscape and resonate with Mason’s determinants of the human stress response ^15^. Our choice of the aversive video paradigm (AVP) as a stress induction method also originates from the need for an experimental protocol that does not involve cognitive engagement, as this project was part of a larger investigation into the influence of stress on working memory and the potential impact of Non-Invasive Brain Stimulation (NIBS) on these effects.

The Aversive video paradigm (AVP) has previously been applied to elicit stress responses in human participants due to the novelty, unpredictability, and uncontrollability of the depicted scenes ^13,14,16^. The clip content of the present study closely resembled that used in previous research ^16,17^. This paradigm is easy to administer and does not require the involvement of more than one experimenter. Moreover, an adequate control condition can be relatively easily established via the introduction of emotionally neutral video clips with otherwise comparable content. Limited research has explored the impact of this stress induction paradigm on brain dynamics using EEG and its specific stress induction-dependent component. Therefore, our study involved a comprehensive analysis, incorporating subjective, physiological, and immunological data collected during exposure to the AVP.

We aimed to document stress-related psychological and physiological alterations induced by this paradigm in line with previous studies. We expected an increase in subjective anxiety as measured by the State-Trait Anxiety Inventory - State (STAI-S), negative emotions as measured by Positive and Negative affect schedule (PANAS), and saliva cortisol levels ^13,16^. We furthermore expected decreased heart rate variability and a deceleration of the heart rate, as reported in previous studies with affective movie stimuli ^18,19^. We expected moreover a stress-related increase of immune activation concerning saliva cytokine levels ^20^. We furthermore expected to observe specific EEG alterations. We hypothesized that stress disrupts cognitive top-down control ^21^ and increases bottom-up processing ^22^, and thus expected respective changes of brain frequency power and connectivity, including a reduction of low frequency oscillations, which are known to be involved in top-down control,^23^ and increased high frequency oscillations, which are involved in bottom-up processing^24^. By analyzing all clips individually we also aimed to observe the dynamics of these effects. By analyzing resting state before and after clip presentation, we aimed to elucidate after effects of stress clips, where we expected to observe increased arousal, and a state of increased anxiety after exposition to the stress-inducing intervention^25^.

## RESULTS

### Subjective Data

The ANOVA conducted for the positive affect scores of the PANAS showed a significant main effect of emotional condition [F (1,77) = 19.29, **p<0.001**, η^2^_p_ = 0.2], a significant main effect of time [F(2.19,168.98) = 55.92, **p<0.001**, η^2^_p_ = 0.42], and significant emotional condition x time interaction effect [F (2.49, 192.14) = 4.606, **p = 0.007**, η^2^_p_ = 0.056]. The post-hoc comparisons (shown in figure 2a.) revealed a decrease of positive affect ratings in both conditions as compared to the first time-point (baseline), with a greater decrease in the stress condition. Post-hoc tests also revealed a decrease of positive affect scores in the stress condition after presentation of the movie clips as compared to control. The ANOVA conducted for the negative affect scores of the PANAS showed significant main effects of emotional condition [F (1, 77) = 109.13, **p<0.001**, η^2^_p_ = 0.586], time [F (2.44, 187.64) = 65.62, **p<0.001**, η^2^_p_ = 0.46], and a significant emotional condition x time interaction [F (2.61, 201.24) = 86.61, **p<0.001**, η^2^_p_ = 0.53]. The post-hoc comparisons (shown in figure 2b.) revealed an increase of negative affect ratings in the stress condition and a decrease in the control condition as compared to the first time-point (baseline). Post-hoc tests also revealed a significant increase of negative affect scores in the stress condition after presentation of the movie clips as compared to the control condition. The ANOVA conducted for the state anxiety scores of the STAI-S showed significant main effects of emotional condition [F (1, 77) = 128.05, **p<0.001**, η^2^_p_ = 0.624], time [F (2.53, 194.57) = 75.625, **p<0.001**, η^2^_p_ = 0.495], and a significant emotional condition x time interaction [F (2.55, 196.27) = 95.09, **p<0.001**, η^2^_p_ = 0.533]. The post-hoc comparisons (shown in figure 2c.) revealed an increase of anxiety scores in the stress condition and no change in the control condition as compared to the first time-point (baseline). Post-hoc tests also revealed a significant increase of anxiety scores in the stress condition after the movie clips as compared to control. Subjective scores thus showed a general increase in anxiety and negative feelings, but a decrease in positive feelings after stress induction. This effect was observed after the start of the clips and persisted until the end of the clips.

**Figure 1.**
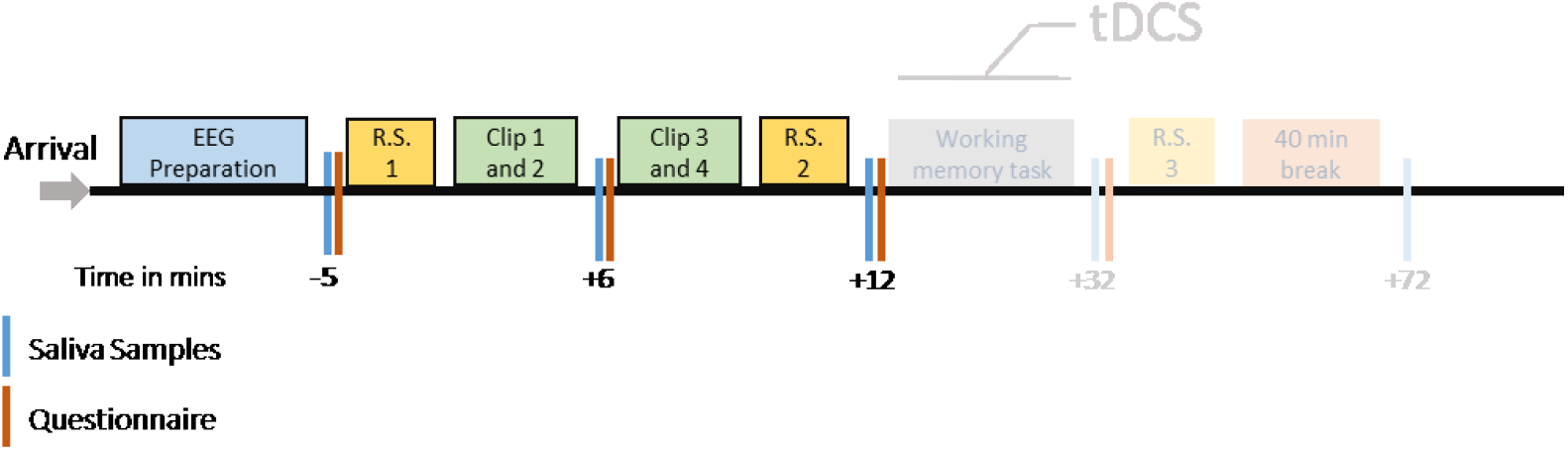
Experimental procedures. After arrival of the participants, the EEG cap was prepared. Then resting state EEG was conducted (R.S.), with 2 min Eyes Open (EO) and 2 min Eyes Closed (EC) EEG. Clips 1-2, and Clips 3-4 denote the movie clips shown (aversive or neutral, according to the control or stress session). Blue bars show the time points of saliva (cortisol, cytokines) sampling and red bars show time points when STAI and PANAS were administered. The time points for saliva sampling and questionnaire conduction in relation to the start of the intervention (clip presentation) are shown below the bar in mins. Heart Rate (using bipolar electrodes positioned on chest) and EEG were recorded during the whole experiment starting from R.S.1 to R.S.3. The procedure was identical for control and stress sessions except for the kind of movie clips shown. The later part of the experiment, including tDCS and working memory task performance, is shown in opaque color and will be reported in detail elsewhere as it is out of scope of the topic of this article.

**Figure 2.**
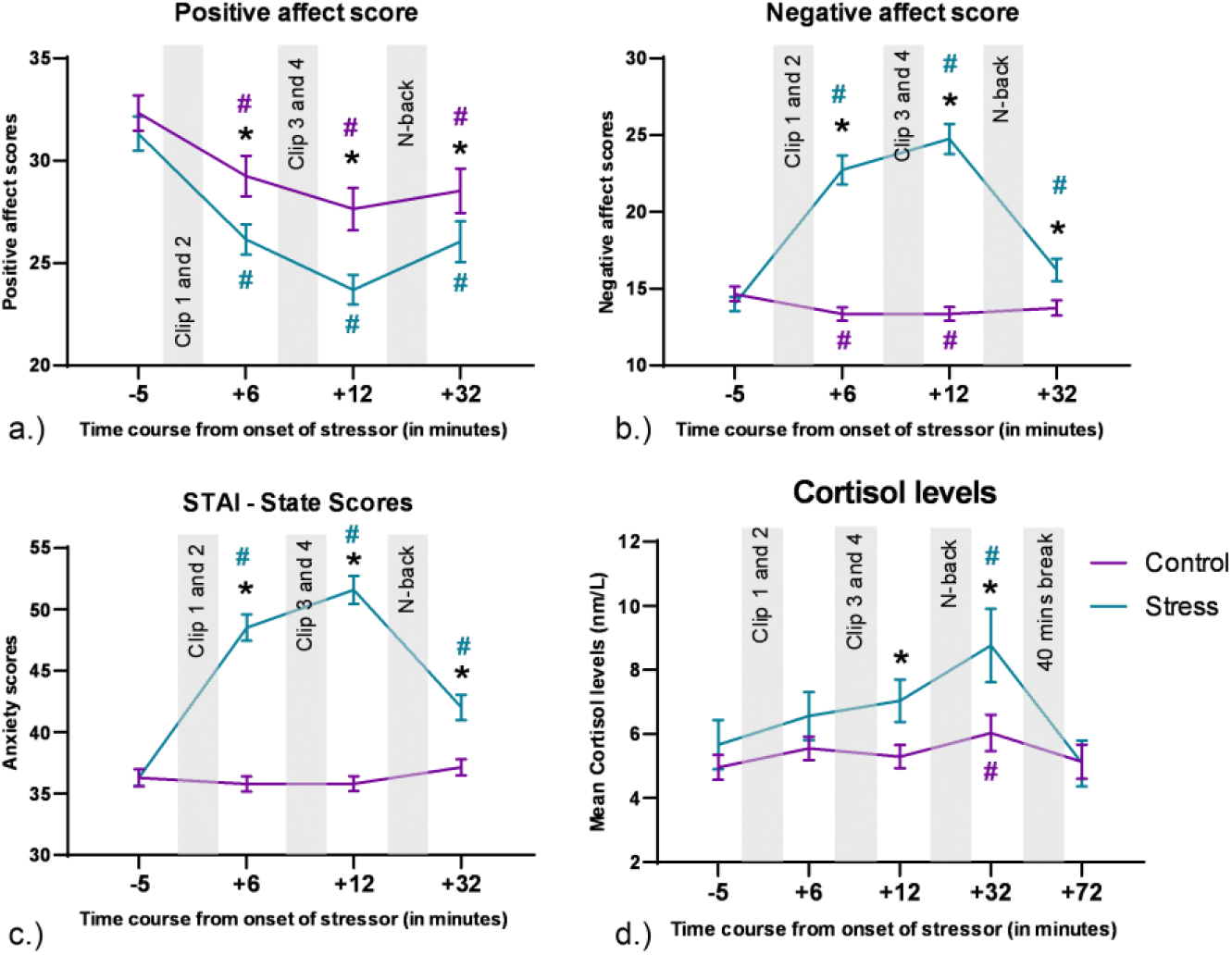
Behavioural and physiological data during stress exposure. a.) Positive affect scores during the experiment. b.) Negative affect scores during the experiment. c.) STAI-S scores during the experiment. For these subjective scores, the y axis denotes the scores in the questionnaire and the x axis shows the time course relative to the start of the movie clips. The last time point of subjective data refers to the time point after conduction of the N-back task, which is out of scope of this paper, and these changes might be also influenced by the task. d.) For cortisol levels, the y axis displays mean cortisol levels (nm/l), and the x axis shows the time course relative to the start of the movie clips. Error bars denote +/-SEM. Asterisks (*) denote significant differences (critical p-value <=0.05) between intervention conditions for each time point. The hash symbol (#) denotes significant differences (critical p-value <=0.05) between time points during intervention and baseline (time point 1) in each condition. The respective color of (#) indicates the respective intervention condition. The last two time points of the cortisol measures were conducted after N-back task performance, which is out of scope of this paper.

### Cortisol

the cortisol analysis revealed significant main effects of emotional condition ([F (1, 69) = 10.25, **p=0.002**, η^2^_p_ = 0.129]) and time point ([F (2.85, 196.81) = 5.893, **p<0.001**, η^2^_p_ = 0.08]), while no significant emotional condition x time interaction was observed [F (2.73, 188.48) = 2.122, p=0.105, η^2^_p_ = 0.03]. Post-hoc tests (shown in figure 2d.) revealed an increase in Cortisol levels for both, control and stress conditions at time point 4 as compared to time point 1 (baseline). Post hoc analyses also revealed a significant increase of cortisol in the stress condition compared to the control condition at time points 3 and 4.

### Cytokines

We analyzed all targeted cytokines via ANOVAs, and for all cytokines, the respective ANOVAs revealed a significant main effect of time, but not of emotional condition, and the respective interactions (for the ANOVA results, refer to table 1). The post hoc analysis conducted for the factor time revealed a common pattern of increased cytokine concentrations in both control and stress conditions during the time course of the experiments. Significant changes of cytokine concentrations relative to baseline in both, control and stress conditions are marked in figure 3.

**Table 1.**
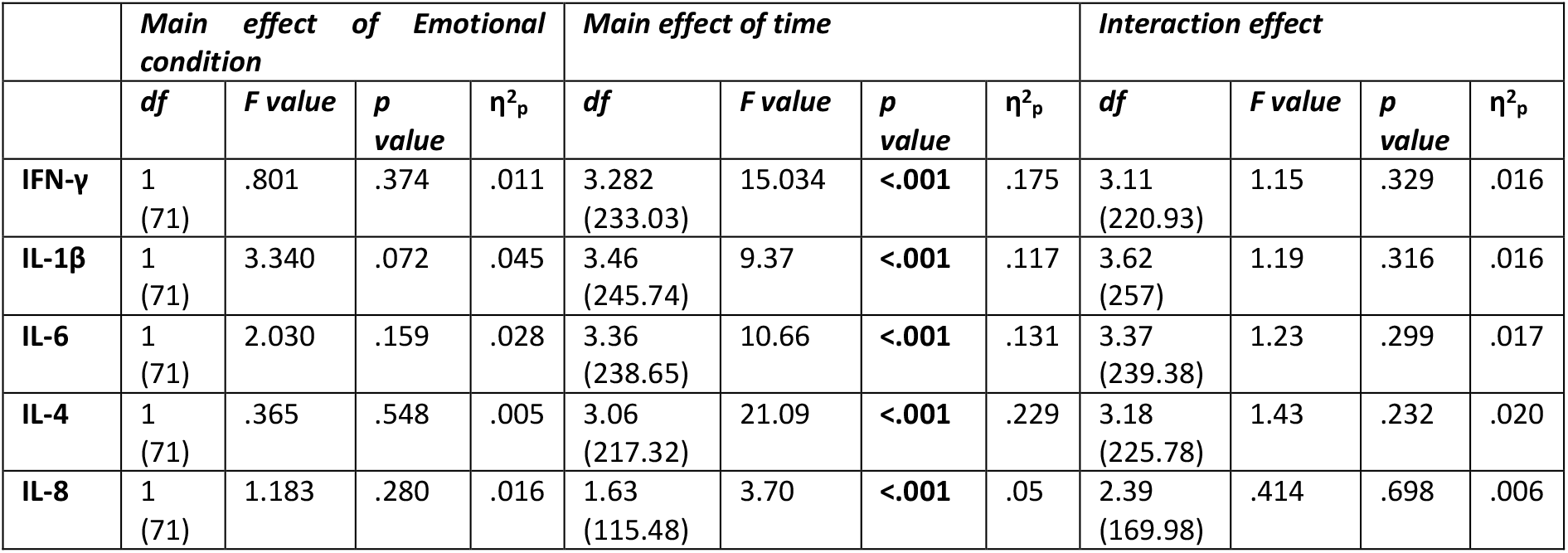
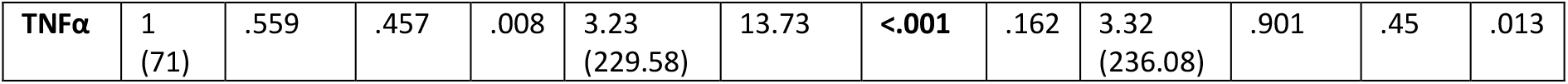
Main and interaction effects of the ANOVAs conducted for all targeted cytokines. Significant p-values are marked in bold.

**Figure 3.**
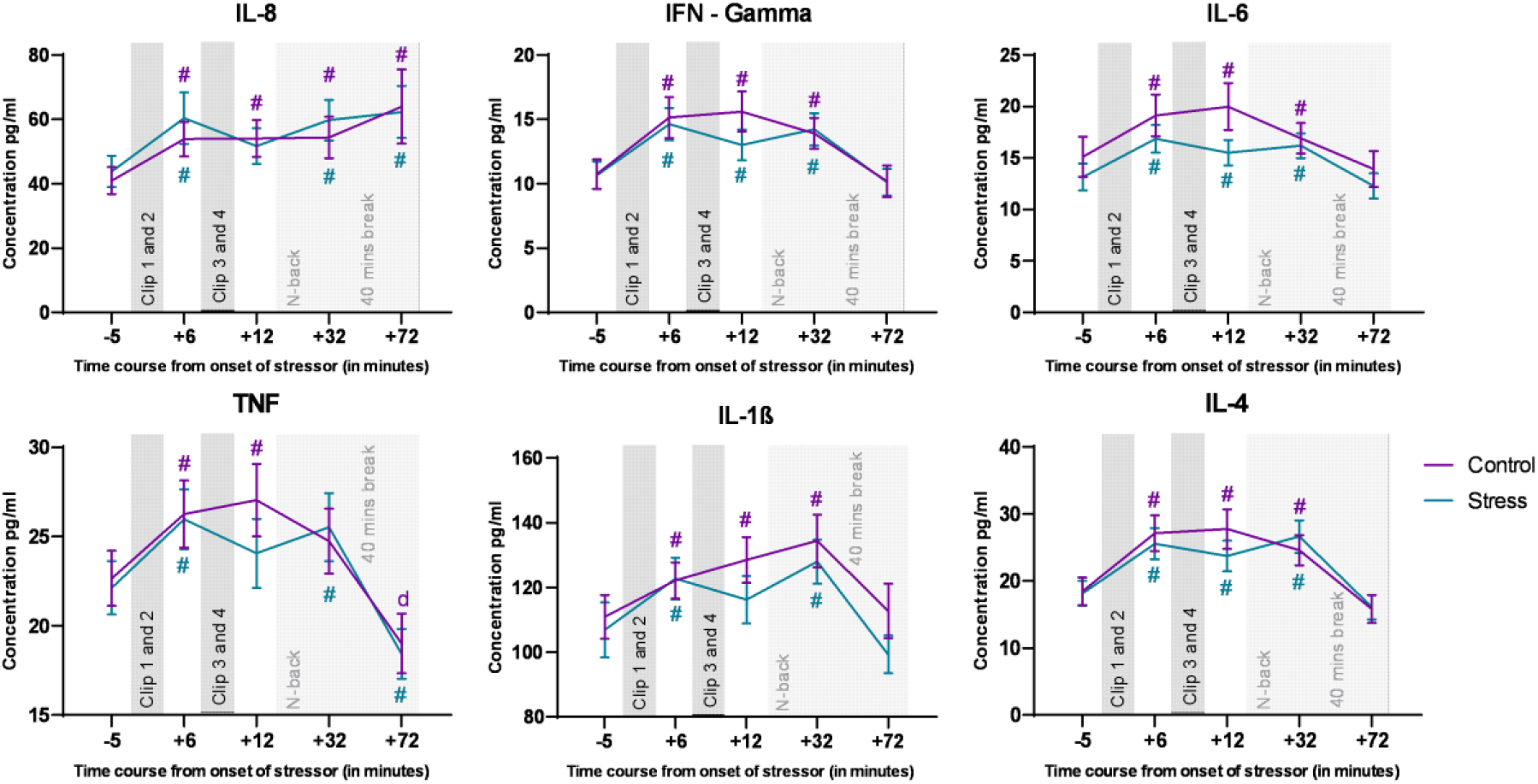
Cytokine levels during the experiment. The Y axis denotes cytokine concentration levels (pg/ml) and the x axis denotes time relative to onset of the movie clips. Hash symbols (#) denote significant differences (critical p-value <=0.05) between baseline (time point 1) and the other time points within each condition. The respective color of (#) denotes the respective condition r. Error bars show +/-SEM. The last two time points were obtained are after N-back task performance, and the dynamics of these time points might be influenced also by the task.

### HR

For heart rate analysis, the ANOVA revealed no significant main effect of emotional condition [F (1, 76) = .132, p=0.718, η^2^_p_ = .002], but a significant main effect of time [F (8.44, 641.67) = 12.63, **p<0.001**, η^2^_p_ = .143], and a significant emotional condition*time interaction [F (7.6, 578.17) = 3.85, **p<0.001**, η^2^_p_ = .048]. Pair-wise post hoc comparisons (shown in figure 4a.) for time points in different emotional conditions revealed a significant increase of HR as compared to baseline (t1) in both conditions during the clips. Post-hoc comparisons between the control and stress conditions for each time point revealed an initial increase of heart rate in the stress condition at the start of clip 1 (t5), and then a deceleration leading to reversal of the difference towards the end of clip 1 (t7). HR differences then remained non-significant between control and stress conditions for the remaining time points.

**Figure 4.**
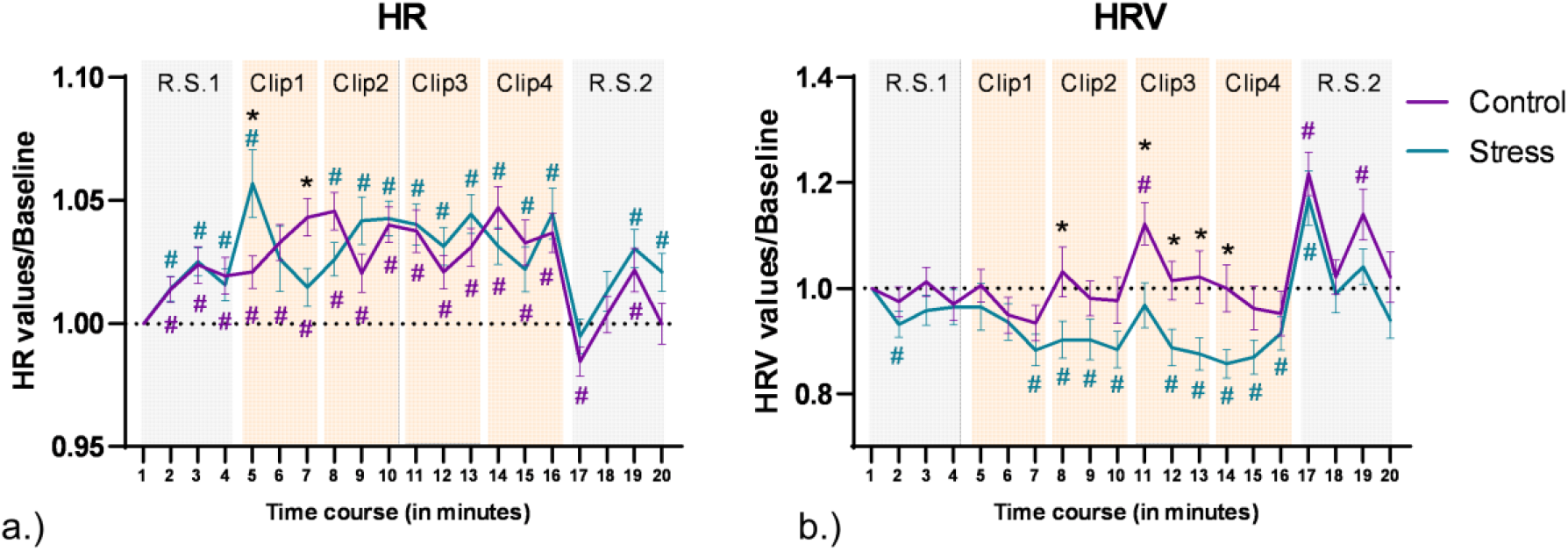
Cardiac data obtained during the experiment. a.) Heart rate during the experiment. The y axis denotes changes of heart rate from baseline, and the x axis denotes each minute of the experiment. Error bars denote +/-SEM. Asterisks (*) denote significant differences (critical p-value <=0.05) between conditions for each time point. The hash symbol (#) denotes significant differences (critical p-value <=0.05) between time points and baseline (time point 1) in each condition. The respective color of (#) denotes the intervention condition. Resting state and clip time points are highlighted in the background with grey and orange colored watermarks (grey represents resting state and orange clips clip presentation). b.) Heart rate variablility during the experiment. HRV was calculated based on RMSSD measures and plotted the same way as heart rate data.

### HRV

For heart rate variability analysis, the ANOVA results show significant main effects of emotional condition [F (1, 76) = 5.412, **p=0.023**, η^2^_p_ = .066], and time [F (9.55, 725.94) = 11.55, **p<0.001**, η^2^_p_ = .132], and a trendwise interaction effect of emotional condition*time [F (9.03, 686.55) = 1.75, **p=0.074**, η^2^_p_ = .023]. The pair-wise post hoc comparisons (shown in figure 4b.) conducted for each time point vs baseline for each intervention condition separately showed a significant decrease of HRV as compared to baseline (t1) in the stress condition (t2, t7-10, t12-17), while in the control condition a significant increase of HRV was observed for single time points (t11, t17, t19). Post-hoc comparisons between control and stress conditions for each time point showed a significantly decreased HRV for the stress condition during Clip 3 (t11-t13), the start of Clip 2 (t8), and Clip 4 (t14).

### Correlations between Stress markers

To test if cytokine responses were related to other stress responses, we conducted Pearson’s bivariate correlations between cytokine area under the curve (AUCi) on the one hand, and cortisol AUCi as well as the AUCi values of HR and HRV on the other for both, control and stress conditions, as in a previous study ^26^. Negative correlations between IL-4 and cortisol (r = -0.233, p = 0.049) and IFN-γ and cortisol (r = -0.262, p = 0.026) emerged in the stress condition only.

### EEG Power analysis

for EEG power calculation during clip presentation, we analyzed all 62 EEG electrodes at different frequency ranges (θ, α, β (low), β (high), γ (low), and γ (high)). We first analyzed each clip individually and later combined the EEG over all clips (clips 1-4). In each analysis we compared the stress clips with the respective control clips. We observed clusters in the Theta, Alpha, low Beta, low Gamma, and high Gamma range. Power was decreased in Theta, Alpha, and low Beta frequencies in each clip and also over all clips combined in the stress condition as compared to the control condition. Alpha power was lower in most of the electrodes, while for Theta the respective clusters of reduced power were situated over frontal, temporal, and occipital regions. For low Beta frequencies, specific clusters of stress-related reduced power were identified in frontal, central, and parietal areas. In the Gamma range, power was significantly larger in clips 2 and 4 for low Gamma, and only in clip 4 for high Gamma oscillations. For both, low and high Gamma, respective clusters of stress-related larger Gamma power were identified over occipital, temporal, and parietal regions. For the exact location and size of these clusters refer to figure 5. Further cluster details for each analysis can be found in the supplementary table 2.

**Figure 5.**
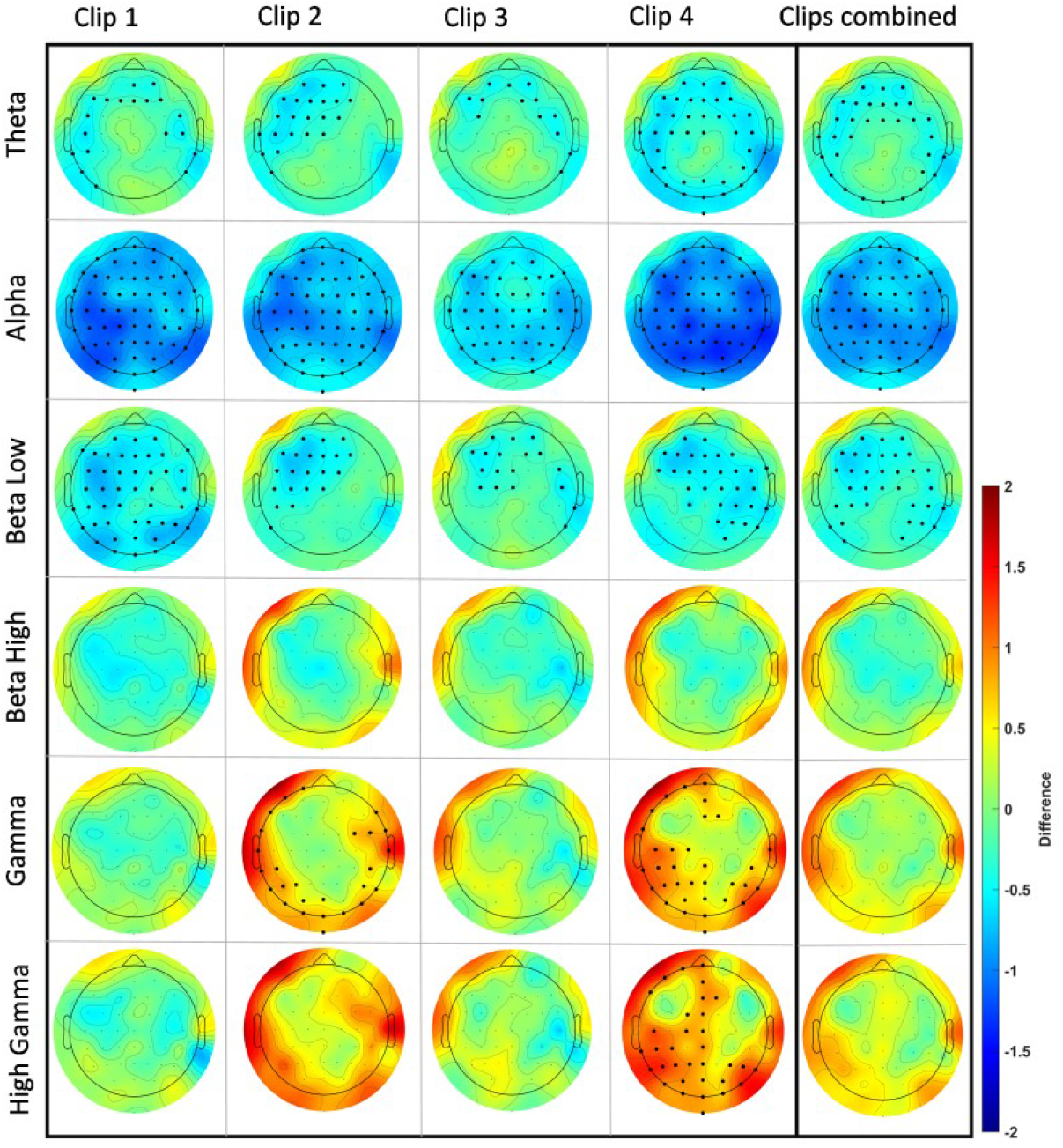
EEG power changes during each clip separately, and for all clips combined for each frequency band of interest. The topographical plots show the mean difference (stress-control) for each block. Significant electrodes in the cluster are marked in bold and were calculated using permutation-based statistics with cluster correction for a critical p-value <=0.01. The color bar indicates the difference value range from 2 to -2. Red shows a higher value and blue a lower value in the stress compared to the control intervention condition.

**Figure 6.**
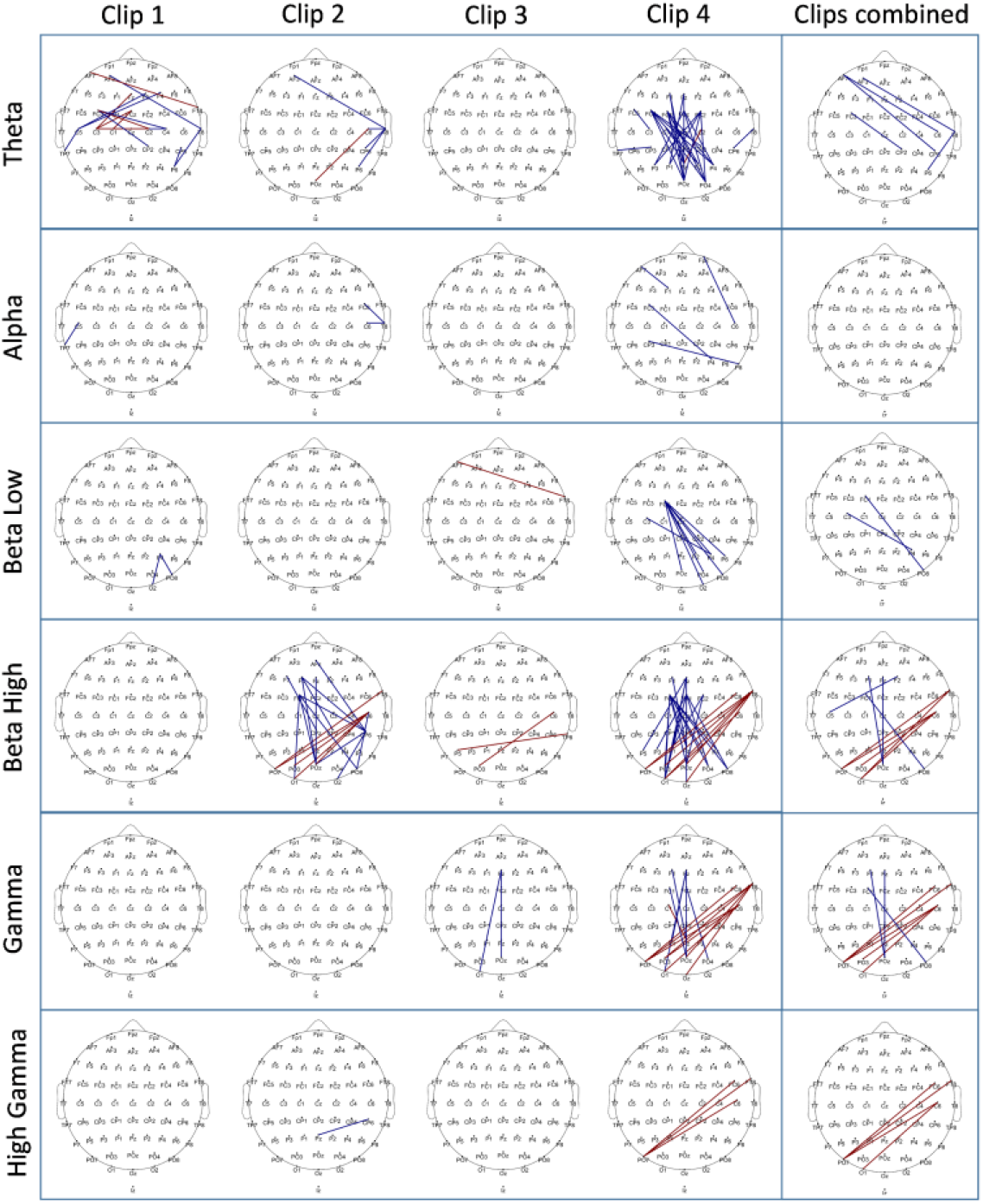
EEG connectivity differences between stress and control conditions shown for each clip separately, and all clips combined for each frequency band of interest. The topographical plots show the significant connections marked in two colors. Blue shows a decrease of connectivity strength and red shows an increase of connectivity strength in stress compared to control conditions. The significance of connections was calculated using permutation-based statistics with maximum statistics based correction which identifies significant connections based on a distribution created by permuted data. The differences were marked as significant for a critical p-value <=0.05.

For EEG power during the resting state for control and stress conditions, we compared resting state 1 (RS1) and 2 (RS2), before and after movie clip exposure for both, control and stress conditions in eyes open (EO) and eyes closed (EC) states for each frequency band. We plotted the topographic difference by comparing RS2 and RS1 (RS2-RS1) for control and stress conditions for both, EO and EC states for each frequency band. We did not identify any significant changes in the control condition for EO and EC states. In the stress condition, we found however significant changes in both, EO and EC states. In the EO state, a significant power increase in high frequency bands, including high Beta, with clusters in parietal and right fronto-temporal areas, low Gamma, and high Gamma, with clusters over almost the whole scalp, was revealed. In the EC state, a significant power increase was identified for all predefined frequency bands with clusters spanning the whole scalp. For the exact location and size of clusters refer to figure 7a, and supplementary table 2.

**Figure 7.**
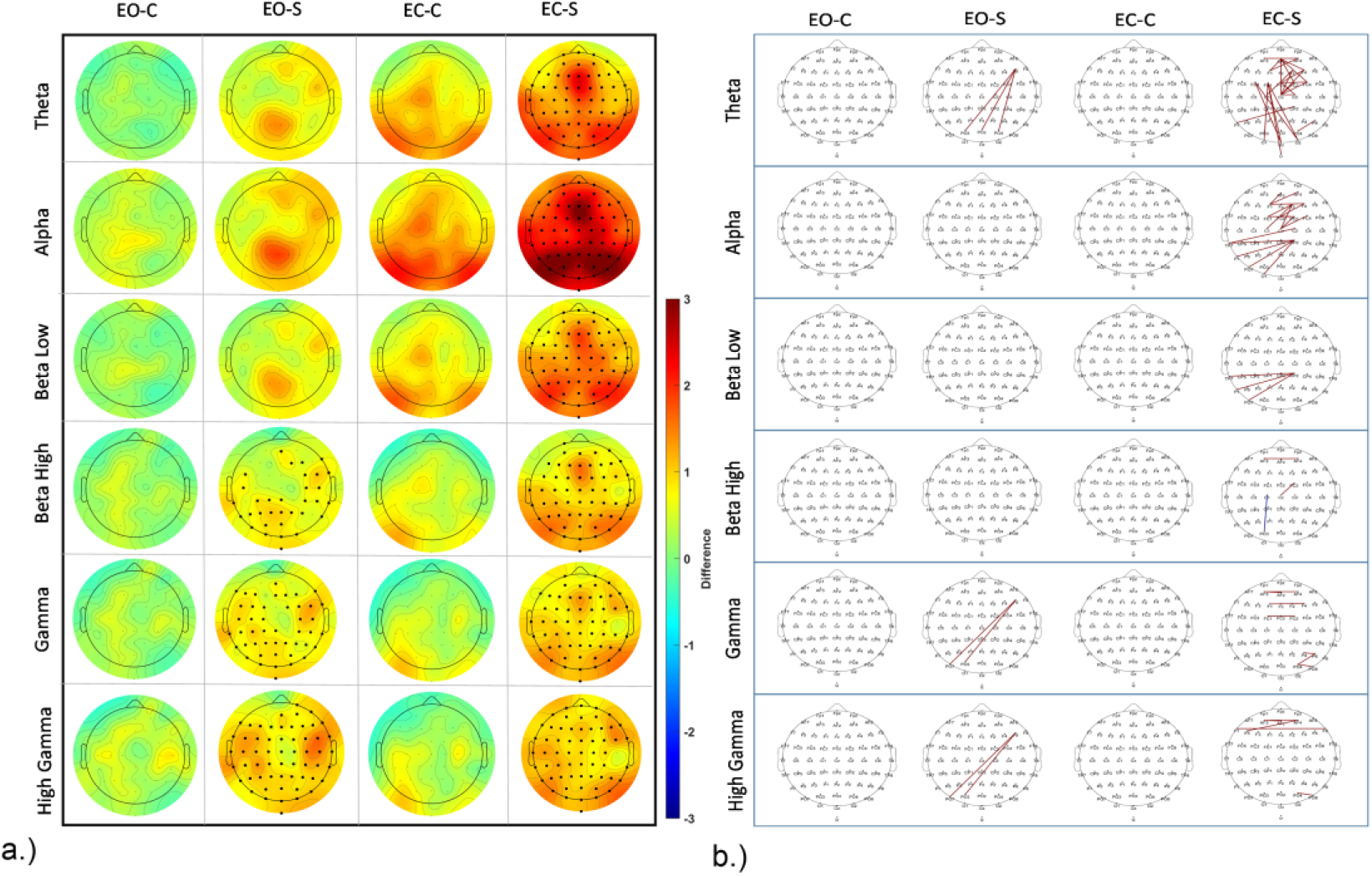
a.) EEG power changes during resting state for both, EO and EC states. The topographical plots show power differences before and after intervention for eyes open and closed conditions, and stress, and control interventions. Significant electrodes are marked in bold. The color bar denotes the difference value range from 3 to -3. Red shows higher power values in the post intervention resting state as compared with pre-intervention, and blue shows reduced power. Statistics were calculated as for the power values during clip presentation. b.) EEG connectivity differences during resting states before and after intervention for both EO and EC states in control and stress conditions. Significant connections are marked in blue and red. Blue indicates a decrease of connectivity strength in the post intervention resting state, and red denotes an increase of connectivity strength in the post-as compared to the pre-intervention resting state. Statistics were calculated as for the power values during clip presentation. EO-C denotes Eyes Open – Control, and EO-S denotes Eyes Open – Stress. EC-C denotes Eyes Closed – Control, and EC-S denotes Eyes Closed – Stress

To analyze group differences during resting states, we compared resting state activity between control and stress conditions during RS1 and RS2 for both, Eyes Open (EO) and Eyes Closed (EC) states for each frequency band. We plotted the topographical difference by comparing stress and control conditions during resting states (stress-control). We found no significant difference between control and stress conditions during RS1 for the EC state, while for EO state, lower power was revealed in the high Beta range in the stress condition, with a cluster over the right frontal, and left parietal regions. For RS2, in the EC state increased power was revealed in the stress condition for high Gamma oscillations. A respective cluster was identified over frontal, temporal and parietal regions. No significant difference between intervention conditions was seen in the EO state in RS2. For exact location and size of all clusters refer to supplementary figure S1 a.

### Connectivity

The connectivity analysis during movie clip presentation was performed for all electrode pairs via the coherence method. The resulting matrix of 62x62 connectivity pairs (excluding self- and mirrored connections) was contrasted between control and stress conditions for each clip, and the resulting significant connections are plotted in figure 6. The analysis was done for all targeted frequency ranges separately, as described for the power analysis. The largest connectivity differences were revealed for Theta, low Beta, high Beta, and low Gamma bands, while only minor differences were present in Alpha and high Gamma frequency bands. We observed the most prominent changes in Clip 4. Here connectivity decreased in the Theta band in the stress vs control condition with respect to fronto–posterior, and fronto-central to posterior connections. In the low Beta frequency band, a relatively decreased connectivity in the stress condition for left fronto-central and right posterior-occipital regions was revealed. In the high Beta and low Gamma frequency bands, similar changes were observed in the stress condition as compared to control, with decreased connections between fronto–posterior regions, but in contrast, increased connections between right temporal and parieto-occipital areas were revealed. Similar changes were revealed in the high Beta frequency band for Clip 2 as well. Please refer to figure 6 for a detailed presentation of all significant connectivity changes (for a list of all connections refer to the supplementary excel file, for an electrode to electrode statistical matrix for each analysis please refer to the supplementary material, Clip Connectivity matrices.

We furthermore performed a connectivity analysis for the resting state as described above for the clip connectivity analysis. We found connectivity alterations in RS2 compared to RS1 (figure 7). We compared RS1 and RS2 for both, control and stress conditions in EO and EC states, as in the respective power analysis above. We did not identify any significant connectivity differences between RS1 and RS2 in the control condition for EO and EC states. For the stress condition, in the EO state we identified significant changes in the Theta, low Gamma, and high Gamma frequency bands between RS1 and RS2. Specifically, an increased connectivity between left frontal (F6) and parieto-occipital regions was observed. In the EC state of the stress condition, significant connectivity increases in the Theta, Alpha, and low Beta frequency ranges emerged, however only few changes were observed in high Beta, low Gamma and high Gamma. Specifically, in the Theta frequency range, a connectivity increase was observed for within-frontal connections and left fronto -occipital connections. In the alpha range, a connectivity increase was significant for within-frontal and posterior-central and left-parieto-occipital connections. In the low Beta frequency range, a connectivity increase was observed only for centro-posterior and left-parieto-occipital connections. In the gamma frequency range, few increased connections within frontal and right parietal regions were observed. Comparing resting state connectivity for control and stress conditions within RS1 and RS2, we found no significant connectivity differences in both EO and EC states. Please refer to figure 7 and S1b (for a list of all connections refer to the supplementary excel file; for an electrode to electrode statistical matrix for each analysis please refer to the supplementary material – Resting state connectivity matrices.

## DISCUSSION

The goal of the present study was to investigate the efficacy of stress induction in healthy human adults through aversive video clips and to explore the dynamics of stress effects on emotion, brain, and body physiology by recording data from multiple modalities. We aimed to replicate previous findings from studies that employed similar stress induction paradigms and to explore previously uncharted territories within the context of this stress induction paradigm ^14^.

Our main results indicate a significant increase of negative affect scores and a reduction of positive affect scores following exposure to aversive video clips, along with increased anxiety scores as measured by the STAI. We also observed elevated salivary cortisol levels, suggesting activation of the hypothalamic-pituitary-adrenal (HPA) axis, in accordance with previous studies using aversive video clips ^13,14,27^. Physiological measurements during the stress condition revealed a decrease of heart rate variability (HRV) during the movie clips as compared to the control condition and an initial increase in HR, which later decelerated again relative to this increase and became similar to HR in the control condition. Uncertainty of events can lead to increased HR ^28^, in accordance with the initial HR increase during the stress condition, as participants were aware about the start of the clips. Previous studies using movie clips as affective stimuli have also shown that movies depicting unpleasant or pleasant stimuli lead to HR deceleration with higher deceleration for aversive clips ^18,19^, which is also in general accordance with the results of the present study, where an initial increase of HR was followed by a relative decrease, indicative for enhanced sensory processing and attention during the clips ^29^. The decrease of HRV during clip presentation in the stress condition is likely caused by reduced parasympathetic activation due to clip presentation and thus a reduced vagal impact on the cardiac rhythm ^30^. Thus, in the present study the clips intended to induce stress led to increased anxiety and negative emotions, increased HPA axis activity, and reduced heart rate variability, and thus psychological, and physiological alterations indicative for stress ^5^.

Furthermore, the study explored the dynamics of cytokine-mediated immune responses to acute stress using the aversive video paradigm. While psychological stress in general is known to influence cytokine activity ^9^, limited research is available for immediate cytokine dynamics following acute stress induction, and specifically no study with AVP, which is a stressor designed to closely resemble real world pure psychological stress, is available. Interestingly, we observed a significant increase of the levels of all explored cytokines during the experiment in both, control and stress conditions. This result is similar to a previous study by Larra et al., who reported similar cytokine enhancements in both, stress and control conditions during the Cold Pressor Test (CPT) ^26^.This increase could have various reasons, including the general design and procedure of the task and stressor, with a novel context in both conditions, which demanded situational involvement. Indeed, previous work has shown that activation of the SNS pathway leads to rapid release of catecholamines that activate nuclear factor κβ (NF-κβ), a transcription factor which triggers the expression of proinflammatory genes and production of cytokines, which is later reduced by the delayed cortisol response ^31,32^. Thus, the secretion of stress-related cytokines is driven by sympathoadrenal activation. For comprehension of the results, it is relevant that the AVP introduces a purely psychological stressor, which does not involve relevant physical or complex cognitive activity, and that the present study involved an active control intervention requiring similar cognitive activity. It thus differs from other stress induction studies reporting cytokine increase, which included stress induction protocols requiring physical or complex cognitive activity, as reported in the review of Slavish and co-workers, who showed that acute stress leads to an increase of inflammatory cytokines, where the most prominent increase was reported in studies comparing an active stress session (especially including a physical stressor) with a rest condition. ^20^ Another relevant aspect is that cytokine analysis with saliva has relevant potential shortcomings, including the impact of oral health, transport to saliva, and the low correlation of immune levels between saliva and blood ^20^. With respect to the association of cytokines and cortisol, we identified similar to Larra et al. a significant negative correlation between cytokine and cortisol levels only in the stress condition, and the cytokines for which we found these correlations were IFN-γ and IL-4, hinting at a possible stressor-specific cortisol-cytokine interaction^26^.

Concerning the effects of the stress induction protocol on oscillatory brain activity, we analyzed absolute power and connectivity differences between stress and control conditions in multiple frequency bands during clip presentation, and in resting states before and after clip presentation. Since stress exerts its effects on a broad range of EEG features, analyzing multiple frequency bands via power and connectivity analyses is required to receive a complete picture ^8^.

In the power analysis conducted for the EEG during clip exposure, we consistently observed a power decrease within the low-frequency range (4-15 Hz) across all stress-inducing clips, but a power increase within the high-frequency range (32-80 Hz) for specific clips. In the Theta range, this reduction was mainly seen in frontal and temporal areas. Alpha reduction, another common EEG alteration during stress, was visible for all electrodes in all clips. In the Beta range, only low Beta (12-15 Hz) showed a decrease over central, frontal, and parietal regions in all clips, while high Beta remained unchanged. In the Gamma and high Gamma frequency ranges, we observed a power increase during presentation of clips 2 and 4, most prominently over occipital, temporal, frontal, and parietal regions.

In the connectivity analysis conducted for the EEG during clip presentation, notable connectivity differences emerged in the Theta, low Beta, high Beta, low Gamma and high Gamma ranges. Connectivity between fronto-central and parietal regions was reduced for all these frequency bands during stress induction compared to the control condition, particularly during Clip 4, indicating a potential influence of clip content. Additionally, during Clip 4 presentation, and when all clips were combined, we observed a connectivity increase between the right temporal and posterior area for high Beta, low Gamma, and high Gamma frequency ranges. In Clip 2, we found such connectivity differences only for the high Beta range.

Comparable alterations of low frequency oscillations during stress exposure have been also reported in previous studies, where stress led to a reduction in frontal theta ^16^, alpha ^33,34^, and low beta ^35^ frequency bands. Low frequency oscillations are known to be involved in top-down processing of information ^23,36^. Previous research has furthermore highlighted a role of gamma activity in negative emotions such as processing negative faces ^37^ and in worry and generalized anxiety ^38^. Increased high frequency power in similar areas as in the present study was also reported in a previous study with aversive affective pictures ^39^. Previous research has moreover suggested an involvement of Gamma range activity in bottom-up processing ^24^.

From a connectivity perspective, frontal to posterior connections are integral for regulatory control, and are the physiological foundation for top-down control of posterior by frontal cortices ^40^. Earlier EEG classifier studies predicting human emotion from movie clip presentation have shown that connectivity in the Beta and Gamma range can reliably predict stress or aversive emotions ^41,42^. This concept finds support in other studies, where specifically disconnection of frontal and posterior areas in the beta range is related to negative emotions ^43,44^, suggesting a suspension of frontal control over emotional or perceptual areas ^44^.

The increase of connections between the right temporal and posterior areas in the present study suggests enhanced processing of aversive emotions during stress exposure, consistent with the known role of the right temporal lobe in processing aversive stimuli ^45^ and the contribution of the parietal lobe to emotion processing ^46^.

Stress is known to lead to loss of top-down control ^21^ and increased bottom-up processing ^22^, which is in accordance with the results our study, which shows a reduction of low frequency power and a reduction of frontal and posterior connections indicative for decreased top-down control. In contrast, the increased high frequency power and connectivity between the right temporal and posterior areas signify an increased bottom-up processing of aversive stimuli.

In the analysis of resting-state power before and after stress induction, a notable difference emerged solely for the stress intervention condition. For the Eyes Open state, an enhancement of high-frequency band power was evident after presentation of the stress clips, particularly in high Beta, low Gamma, and high Gamma bands. In the Eyes Closed state, a significant power increase spanned all frequency bands of interest. Comparing resting-state EEG power for control and stress conditions after presentation of the movie clips unveiled selectively enhanced High Gamma activity in the stress group.

No connectivity differences were observed between stress and control clips in both resting states. However, after presentation of the stress-inducing clips in the Eyes Closed state, increased Theta connectivity in the right frontal area and between left fronto-central and occipital regions was observed as compared to pre-intervention resting state (baseline). Moreover, in the Alpha frequency range, connectivity rose within the frontal area and between parietal and left temporal regions as compared to baseline.

Increased power in all frequency bands during resting state has been reported during enhanced uncertainty and anxiety ^25^ and a rise in high frequency power (Beta, Gamma) during resting state has also been associated with increased worry and arousal ^38,47^. Moreover, enhanced low frequency (Theta and Alpha) connectivity in resting states has been reported in dysphoria patients associated with enhanced rumination and self-focus ^48^. Thus, the results of the present study are in accordance with a state of enhanced cogitation and arousal after aversive video clip presentation, which was not observed after the neutral clips.

Our study aimed to assess stress induction in healthy adults using aversive video clips and investigate its impact on emotion, physiology, and brain activity. Exposure to aversive clips resulted in increased negative affect, reduced positive affect, and elevated anxiety scores. Physiologically, salivary cortisol levels rose, and altered heart rate dynamics indicated enhanced stress. The cytokine analysis showed increased levels during both stress and control conditions, suggesting an unspecific immune response by conduction of the study. The EEG analyses revealed reductions of low-frequency power, increased high-frequency power, and altered connectivity patterns during stress induction, indicating potential disruptions of top-down control and activation of bottom-up processing. Resting-state EEG showed increased power across all frequency bands after stress induction, suggesting sustained effects of stress induction on neural activity even after stressor presentation. Our findings provide a comprehensive understanding of the intricate interplay between psychological stress, physiological responses, and neural dynamics in healthy adults, highlighting the multi-faceted nature of the stress response.

### Limitations and Future Directions

While this study identified clear psychological, and physiological effects of the AVP protocol indicative for its suitability as a stress-inducing tool, some limitations should be taken into account. The stress induction procedure used in this study includes factors beyond pure stress, such as disgust, potentially influencing the observed responses. The consistent clip sequence, while maintaining paradigm continuity, introduces potential confounding effects from specific clip content interactions with neural dynamics. Additionally, the inclusion of only males limits the generalizability of findings across sexes. To address these limitations, future studies should diversify participant populations, including both sexes. Randomized clip orders would help to disentangle unspecific content effects and better isolate stress responses. The conduction of a working memory task after the clips also raises the possibility of interferences between task-induced anxiety and clip effects for the data obtained after clip presentation, warranting further investigation in future research.

## Conclusions

This study revealed the impact of aversive video clips on subjective, physiological, and brain dynamics observed via EEG. Elevated negative affect, anxiety, and cortisol levels as well as diminished HRV are indicative for increased stress during the paradigm. Loss of top-down control and increased bottom-up processing is suggested by EEG data alterations, thus proposing specific alterations of brain function due to increased stress and negative emotions. Our results paint a multi-modal landscape of stress effects caused by this paradigm which might be advantageous for future stress-related research due to its ability to target only psychological aspects of stress.

## Methods

### Participants

Seventy-eight healthy, right-handed male non-smokers aged between 18 and 40 years participated in the study (M = 25, SD = ±4.16). A medical check before the experiment made sure that the participants did not suffer from any chronic or acute disease, did not have any psychiatric or neurological disorder, did not suffer from any inflammation of the mouth, were not taking any central nervous system (CNS) acting medication, did not work at night or changing shifts during the timeframe of the experiment, nor had any implants or devices in their body. Participants with a history of epilepsy or traumatic brain injury were also excluded. In a pre-experimental phone interview, it was made sure that the participants did not have any major physical or emotional trauma, and that they had no habit of watching extremely violent movies. The study was approved by the Ethics committee of the Leibniz Research Centre for Working Environment and Human Factors at TU Dortmund (IfADo) and aligns with the Declaration of Helsinki. Participants were informed about their right to withdraw from the experiment at any time.

### Stress Induction Procedure

Stress was induced using aversive video clips from the movie “Irreversible” by Gaspar Noe, while control condition clips were extracted from the movie “Comment j’ai tué mon père” by A. Fontaine. Each clip had a duration of 3 minutes, with four clips presented in a predetermined sequence. Control and stress condition clips were carefully matched for duration, luminance, and presence of humans, differing only in violence levels. The first two stress clips depicted violence against a female, while the last two depicted violence against a male. The first two control clips involved a party scene, and the last two clips involved long shots of people walking.

Both movies were in French language without subtitles. Only two of the included participants had French language proficiency. Before viewing, participants received a brief introductory text instructing them to watch the movie clips attentively from an eye-witness perspective.

### Subjective Data

Two questionnaires were administered at various time points during the experiment: the Positive and Negative Affect Schedule (PANAS) ^49,50^ and the State-Trait Anxiety Inventory-State (STAI-S) ^51,52^. The PANAS assessed positive and negative feelings and included 20 questions, which had to be rated on a 5-point scale ranging from 1 (not at all) to 5 (very much), while the STAI-S evaluated state anxiety levels, also included 20 questions which had to be rated on a scale ranging from 1 (not at all) to 4 (very much). Participants were given the choice to fill in the questionnaires in either German or English language based on their personal preference. Data were collected using the online Sosci survey platform ^53^ and subsequently analyzed using custom MATLAB scripts.

### Heart Rate (HR) and Heart Rate Variability (HRV) Data

HR was continuously monitored during the experiment using two bipolar electrodes connected to Bipolar inputs of a NeurOne JackBox (Neurone, Finland)— one electrode was attached below the right clavicle and the other electrode was attached above and left to the umbilicus. HR data were sampled at 2000 Hz and analyzed using HEPLAB ^54^, an EEGLAB plugin ^55^, along with custom-made MATLAB scripts. Inter-beat intervals (IBIs) were calculated, and HR was computed for each 1-minute interval during the experiment during resting states and presentation of the movie clips (for details, refer to figure 1). HRV was determined using the Root Mean Square of Successive Differences (RMSSD) method ^56^ via a custom-made MATLAB script for identical 1-minute bins as for HR.

### Cortisol

Saliva samples were collected at various time points (for exact time points and intervals refer to Figure 1) during the experiment using Sarstedt Salivettes. Cortisol levels were analyzed using the Cortisol Saliva ELISA (TECAN/IBL International) with following the manufacturer’s instructions. Absorbance at 450nm was measured using a GloMax^®^ Multimode Microplate Reader System (Promega). Cortisol data from only 70 participants were analyzed due to the limited availability of kits and insufficient saliva samples from some participants.

### Cytokines

Cytokine levels in saliva were measured twice per sample using the LEGENDplex Human Essential Immune Response panel (BioLegend), as described in previous studies ^26,57^. Briefly, 10 μl saliva or standard control solution was added to a V-bottom 96 well plate mixed with 30 μl assay buffer and 10 μl beads and incubated for 2 hours at room temperature (RT) on a shaker at 800 rpm. After washing with 200 μl wash buffer (provided in the kit), a 10 μl biotinylated detection antibody mix was added to the beads and incubated for 1 hour at room temperature on a shaker at 800 rpm. Then, 10 μl PE-conjugated streptavidin was added, followed by an additional incubation for 30 min at RT on the shaker at 800 rpm. After two washes with wash buffer, the PE fluorescence intensity of the beads was measured on an LSRFortessa flow cytometer (BD Biosciences). Bead populations were identified by FSc/SSc features and fluorescence intensity in the APC channel. Approx. 200 beads per analyte were acquired. Data were analyzed using the LEGENDplexTM Data Analysis Software (VigeneTech). All samples from a participant were analyzed on the same day to avoid inter-assay variation. Cytokines targeted for analysis were Interferon-gamma (IFN-γ), Interleukin – 1Beta (IL-1β), Interleukin – 6 (IL-6), Interleukin – 4 (IL-4), Interleukin – 8 (IL-8), and Tumor necrosis factor - alpha (TNFα). We selected this profile of six essential cytokines to obtain a broad overview of stress effects on cytokines by targeting a mix of pro-inflammatory (IFN-γ, IL8, IL-1β, TNFα, and IL-6) and anti-inflammatory (IL-4) cytokines. These cytokines have been obtained in previous studies exploring the effects of stress on cytokines ^20,26,57^ . Cytokine data from 70 participants were analyzed due to the reasons given above.

### EEG data Acquisition and processing

EEG was recorded during resting states (RS) and during movie clip presentation against a reference electrode placed over the left mastoid, using sintered Ag-AgCl electrodes at 62 positions in accordance with the international 10-20 EEG System (NeurOne, Finland). Electrode impedance was monitored and kept below 10 kOhm throughout the experiment. The sampling rate was 2000 Hz at an analog-to-digital precision of 24 bits. Offline EEG analysis was done using EEGLAB (v2022.1) ^55^, Brain Vision Analyzer (Version 2.2.0, Brain Products GmbH, Gilching, Germany), Fieldtrip ^58^, and custom MATLAB commands.

Data preprocessing was identical for all datasets, involving filtering (1-80 Hz) using the EEGLAB *‘pop_eegfiltnew’* command. Automatic rejection of noisy channels using EEGLAB’s clean raw data plugin to remove channels with stimulation electrodes and other noisy channels was performed. The criterion given to the function was – ‘Flatlinecriterion’ – 5, ‘Channelcriterion’ – 0.65, and ‘Linenoisecriterion’ – 5. Data were then average-referenced using the *‘pop_reref’* EEGLAB command. To remove 50Hz line noise we used the Cleanline plugin of EEGLAB ^59^. Later, eye and muscle artifacts were removed using the artifact removal plugin in EEGLAB, which uses blind source separation (BSS) techniques to remove muscle and eye activity ^60,61^. Later, the data were epoched (for both, clips and RS) into 2-second segments with no baseline. The removed channels were then interpolated using the spline interpolation method in EEGLAB. We then ran an ICA decomposition using the EEGLAB ‘runICA’ command. The resulting decompositions were labelled using the IClabel plugin ^62^. Decompositions having more than 70% probability to be eye, muscle, line noise, or channel noise artifacts were removed from the data.

### Power Analysis

The power analysis employed the EEGLAB ‘STUDY’ function ^55^ and custom MATLAB scripts. The *spectopo* function was used for power estimation with a window size of 998ms and overlap of 495ms, and spectra were generated for all 62 channels and averaged within six frequency bands (θ – 4 to 8 Hz, α – 8 to 13 Hz, β (low) – 13 to 15 Hz, β (high) – 23 -32 Hz, γ (low) – 32 to 50 Hz, and γ (high) – 50-80 Hz). *Spectopo* uses the Pwelch method of the MATLAB signal processing toolbox for power estimation with overlapping hamming windows. The analysis was performed for all 62 electrodes and significant electrodes were identified for each analysis using Montecarlo-based permutation statistics with cluster correction (see data analysis). Topographical plots were generated using EEGLAB’s *topoplot* function.

### Connectivity

Connectivity analysis was conducted using Fieldtrip ^58^. Preprocessed data from EEGLAB were imported using the *‘ft_preprocessing’* command, frequency decomposition using the *‘ft_freqanalysis’* command with the following input parameters [method = mtmfft, taper = dpss, output = fourier, foilim = [2 80], tapsmofrq = 2], and analyzed for connectivity by the *‘ft_connectivityanalysis’* using the coherence (coh) method.

Coherence as a measure for assessing functional connectivity between EEG signals is the most commonly used method to calculate linear dependencies between two channels in the frequency domain. This method was employed as it does not need any a priori hypothesis and previous research has shown that coherence as a measure can reliably differentiate stress indices as compared to phase lag measures ^63^.

Coherence is defined as ratio of cross-spectra to the product of auto-spectra of two signals in a specific frequency band ^64^-

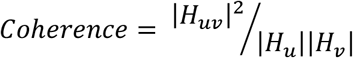

where |*H*_*uv*_|^2^ is the cross-spectrum between two signals, and |*H*_*u*_|, |*H*_*v*_| are the auto-spectra of individual signals. The coherence values range between 0 and 1, where 0 indicates no linear coupling and 1 indicates maximum linear coupling. Coherence was calculated between all channels (62x62), for each epoch, in the six frequency bands introduced above. Significant connections were identified for each analysis using Montecarlo-based permutation statistics with the maximum correction method (max correction) implemented in Fieldtrip, and topographical plots were generated using the EEGLAB *topoplot* function.

### Experimental Procedure

Participants underwent a three-session experimental protocol, starting with a practice session aimed at reducing unspecific experiment- and context-related stress, conducting a medical check, and training of the participants in a working memory task, which will be reported in detail in another publication. Subsequently, two experimental sessions—control and stress conditions—were administered, with randomized and counterbalanced session order. All experiments were conducted between 12 a.m. and 6 p.m. to control for endogenous cortisol activity.

For the experimental sessions, participants arrived one hour prior to EEG scanning for electrode placement. The experiment began with baseline cortisol and questionnaire assessments, followed by an EEG resting state recording (2 minutes with eyes open and 2 minutes with eyes closed). Subsequently, the movie clips were presented on a monitor, with 30 cm eye to screen distance and audio output provided via standard in-ear earphones. Saliva samples were obtained before starting the experiment as baseline and then after the first two movie clips, and then the remaining two clips were shown. Afterwards, another set of saliva samples and questionnaires were collected, followed by the final resting state EEG recording, as described above. The complete experiment included two parts: the first part is described above, and a second part, in which working memory performance was assessed via an n-back task combined with non-invasive brain stimulation, will be described elsewhere. For an overview, please refer to figure 1.

### Data Analysis

Data analysis based on ANOVAs was performed using SPSS version 29.0.0 (IBM Corp., Armonk, New York, USA). We performed Repeated Measure (RM) ANOVAs for subjective data, cortisol, and cytokines. The emotional intervention condition (with the levels control, and stress), and time (t1, t2, t3, t4, t5) served as within subject factors. RM ANOVAs were also performed for HR and HRV data with the emotional intervention condition (control, stress), and time bins (t1-t20) as within subject factors. Sphericity was tested for all ANOVAs with the Mauchly test, and Greenhouse-Geisser corrections were applied when appropriate. For post-hoc tests, Fisher’s Least Significant Difference (LSD) test was used. The critical alpha level was set at 0.05 for all tests. Correlation analyses were performed via Pearson correlations. For correlation analyses, the area under the curve (AUC) was calculated for cytokines, cortisol, HR, and HRV according to the formula ^65^ –

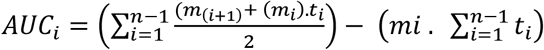

where m_i_ denotes single measurements, t_i_ denotes the time distance between the measurements, and n denotes the total number of measurements. Correlations were then calculated based on these AUC values, similar to a previous study ^26^.

EEG data were analyzed and plotted using the open-source EEG data analysis software EEGLAB and Fieldtrip ^55,58^ implemented in MATLAB (R2020b), as described above. Statistical testing for EEG power involved Monte-Carlo based permutation testing with cluster correction, implemented in the Fieldtrip toolbox in EEGLAB, with the number of permutations set to 8000. Cluster-based test statistics were calculated by comparing each data point using t-tests, and t-values which were significant at an alpha level of </= 0.01 were clustered together based on their spatial and spectral adjacency. The clusters were then aggregated, and their t-value sums used to identify the maximum cluster-level statistics. Significant clusters were marked. For connectivity analysis, we used a similar permutation testing method with a maximum correction method with a critical alpha level of </= 0.05 to control for familywise error rate ^55,58,66^

## Acknowledgments

This work was funded by DAAD (Deutscher Akademischer Austauschdienst) GSSP (Graduate School Scholarship Programme) grant to S.R.

## Author Contributions

S.R., Y.F., M.M.S., and M.A.N. designed the experiment. S.R. and N.M. collected the data. S.R. analyzed behavior, physiological, and EEG data. M.C. analyzed immunological data. S.R. prepared the figures. S.R. wrote the main manuscript text. T.K., and M.A.N. critically revised the manuscript and provided expert comments.

## Data Availability Statement

Required data can be made available upon reasonable request to the corresponding author.

