## Supplementary material for "Multimodal assessment of acute stress dynamics using an Aversive Video Paradigm (AVP)": Clip Connectivity Matrices.pdf

### Theta Range Connectivity Matrix (Clip 1 – 4; Clips Combined)

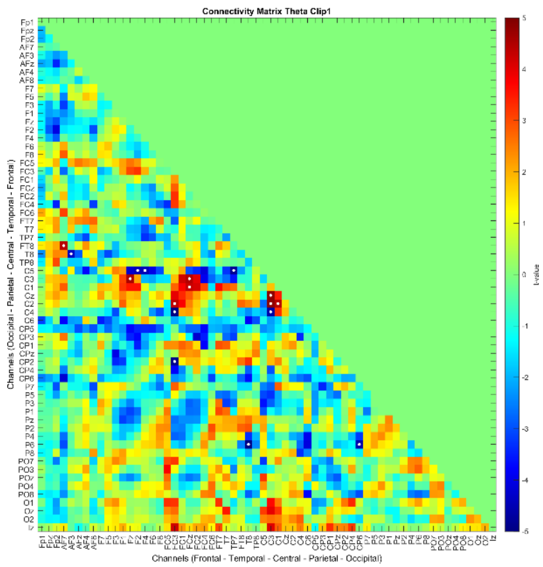


Clip 1

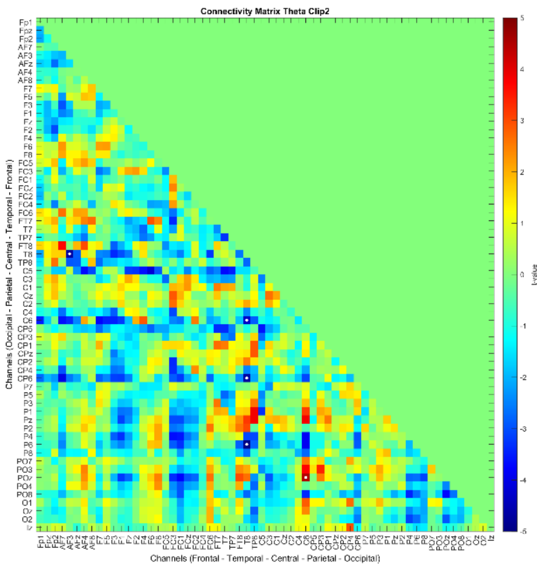


Clip 2

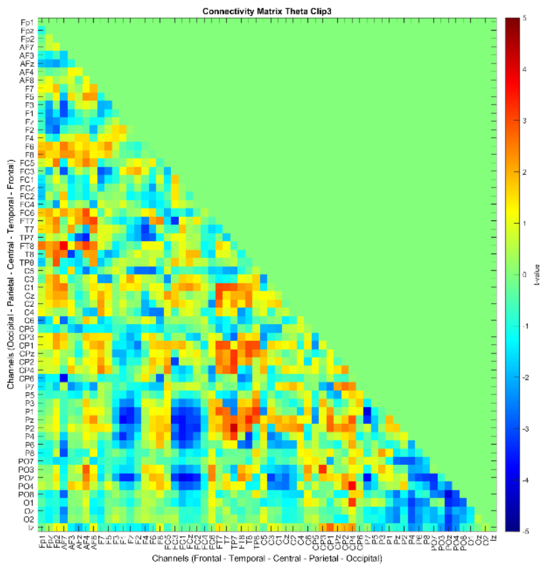


Clip 3

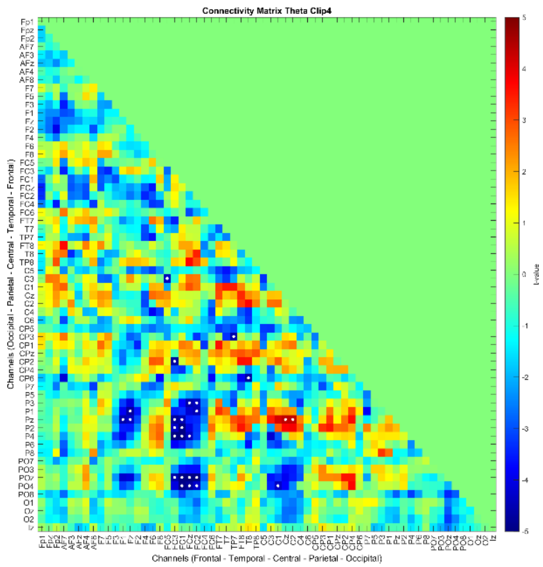


Clip 4

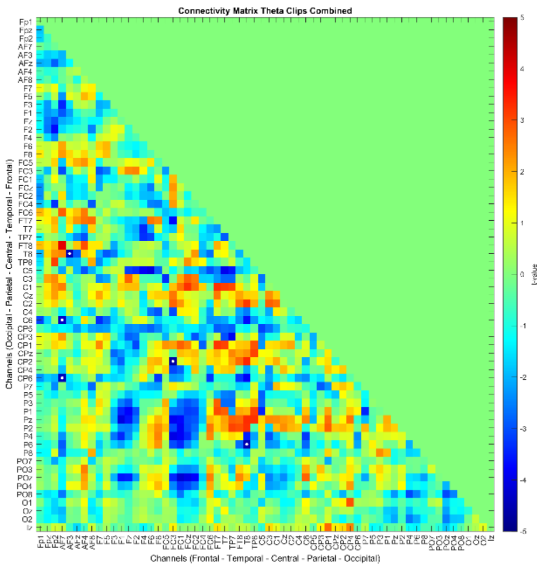


Clips Combined

Analysis was done between control and stress conditions for each clip separately and for the combined clips. The color of connections denotes the direction of connectivity alterations, where red denotes increased connectivity in the stress condition and blue reduced connectivity compared to the control condition. Significant points are highlighted with a white dot. The order of channels is Frontal - Temporal - Central - Parietal - Occipital.

### Alpha Range Connectivity Matrix (Clip 1 – 4; Clips Combined)

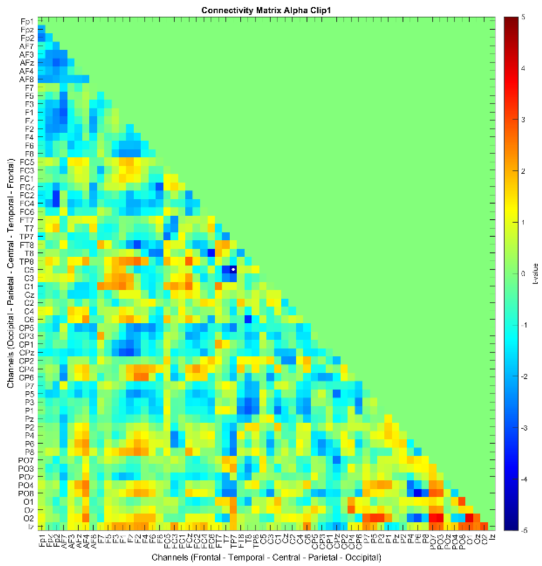


Clip 1

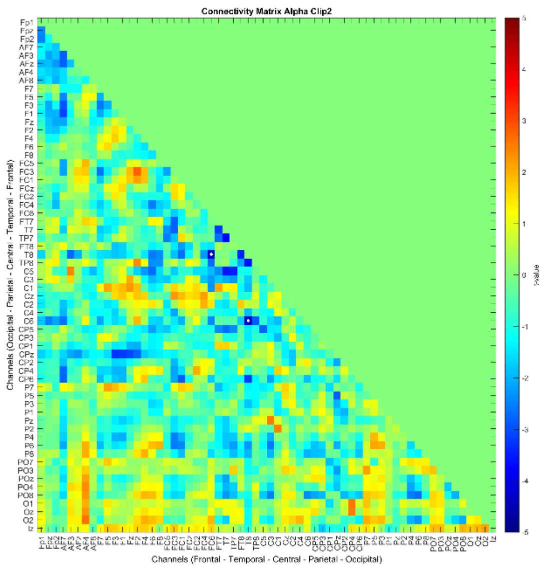


Clip 2

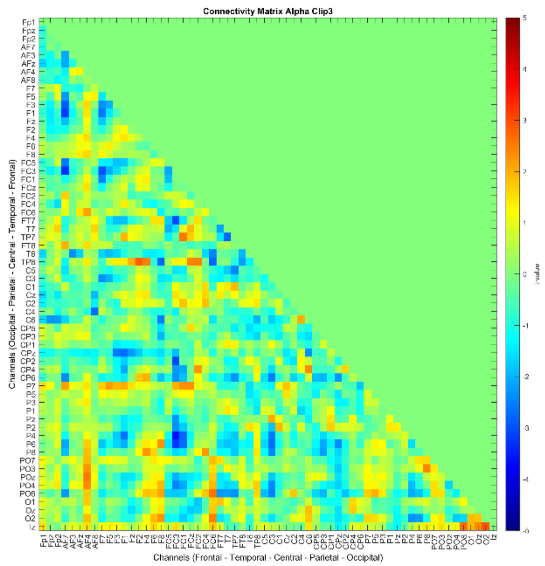


Clip 3

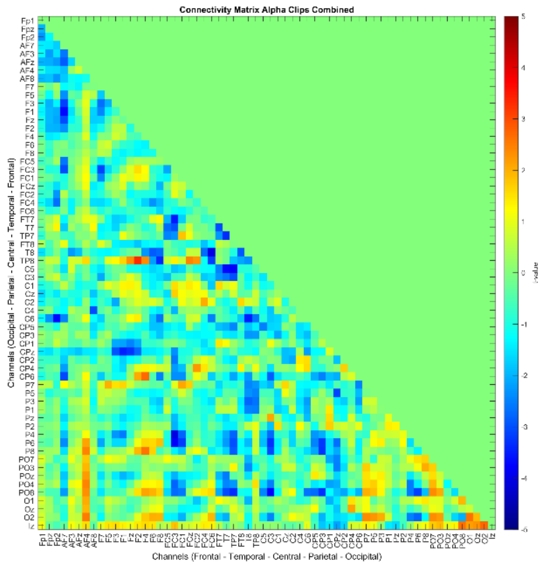


Clip 4

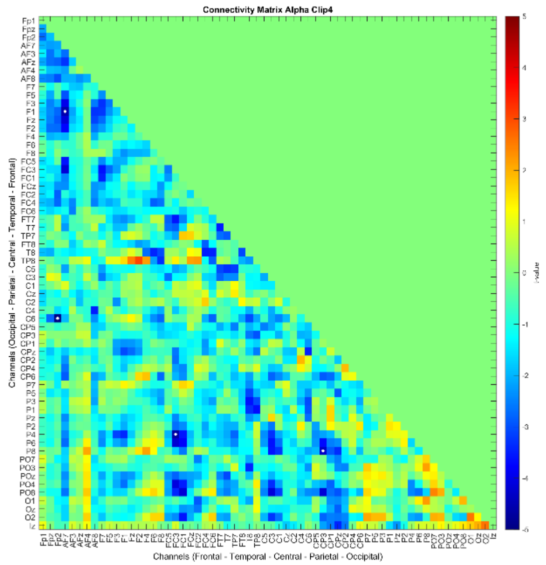


Clips Combined

Analysis was done between control and stress conditions for each clip separately and for the combined clips. The color of connections denotes the direction of connectivity alterations, where red denotes increased connectivity in the stress condition and blue reduced connectivity compared to the control condition. Significant points are highlighted with a white dot. The order of channels is Frontal - Temporal - Central - Parietal - Occipital.

### Low Beta Range Connectivity Matrix (Clip 1 – 4; Clips Combined)

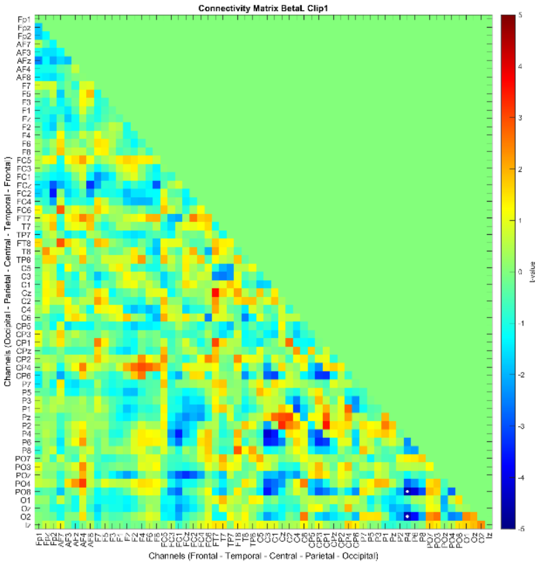


Clip 1

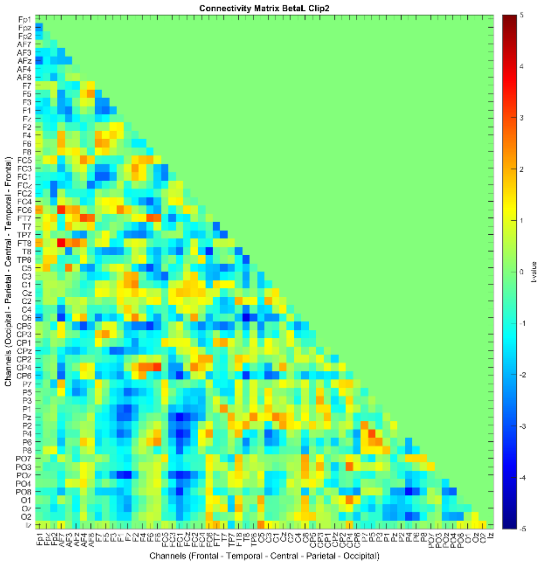


Clip 2

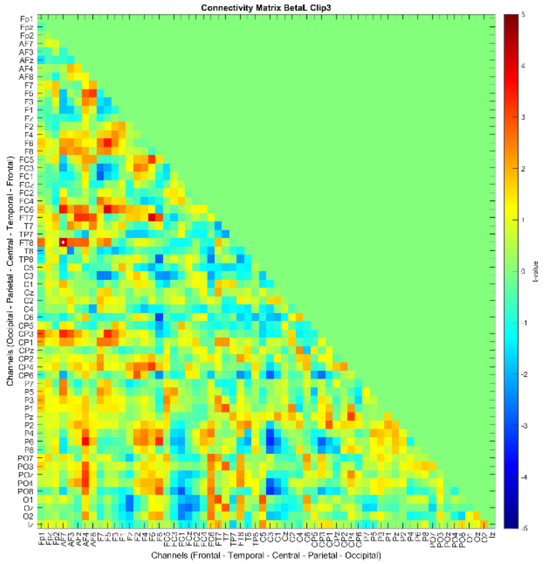


Clip 3

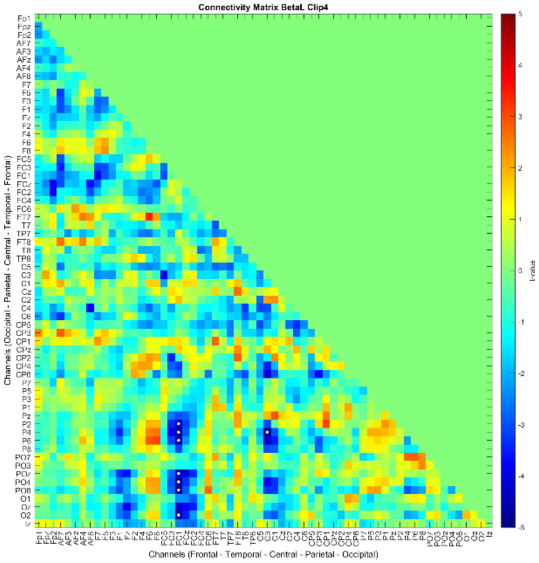


Clip 4

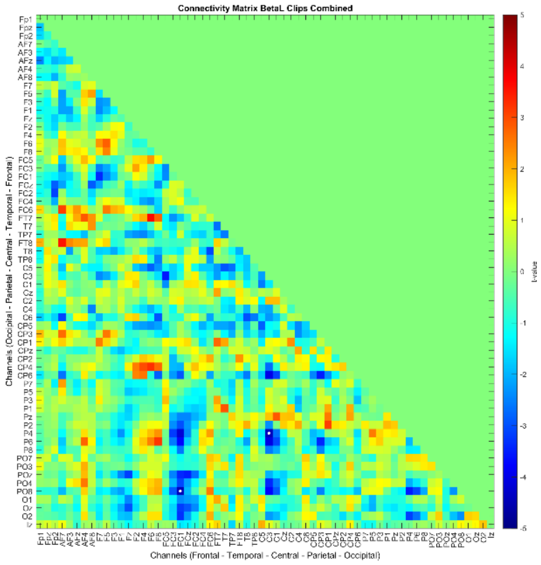


Clips Combined

Analysis was done between control and stress conditions for each clip separately and for the combined clips. The color of connections denotes the direction of connectivity alterations, where red denotes increased connectivity in the stress condition and blue reduced connectivity compared to the control condition. Significant points are highlighted with a white dot. The order of channels is Frontal - Temporal - Central - Parietal - Occipital.

### High Beta Range Connectivity Matrix (Clip 1 – 4; Clips Combined)

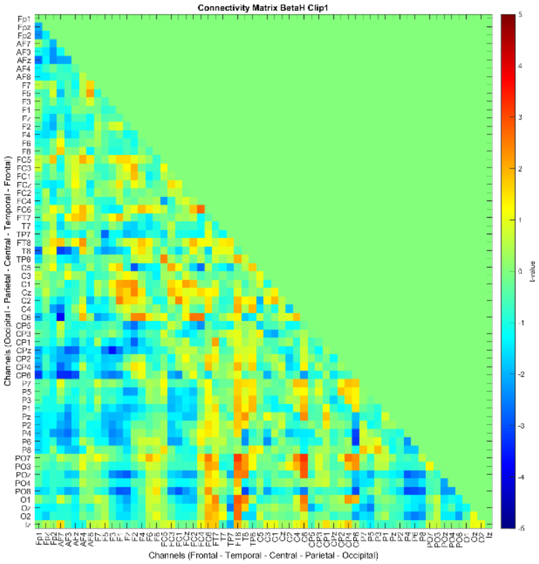


Clip 1

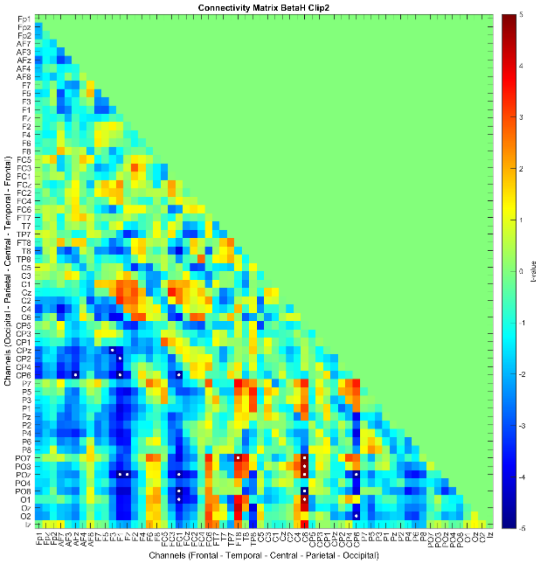


Clip 2

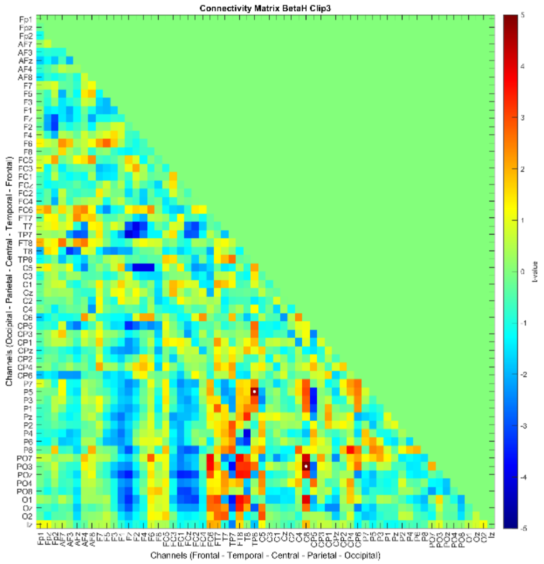


Clip 3

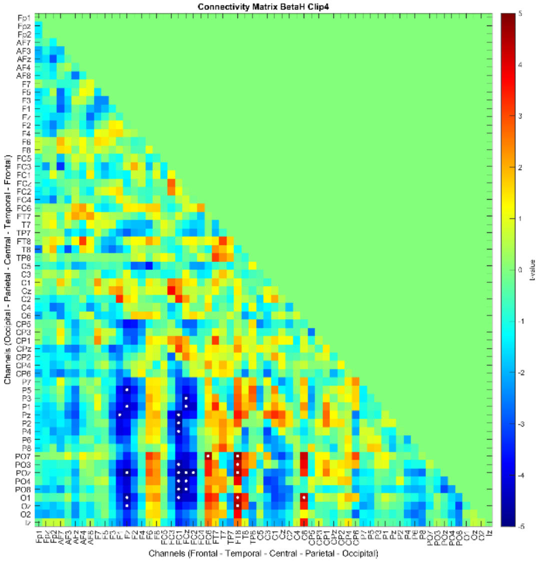


Clip 4

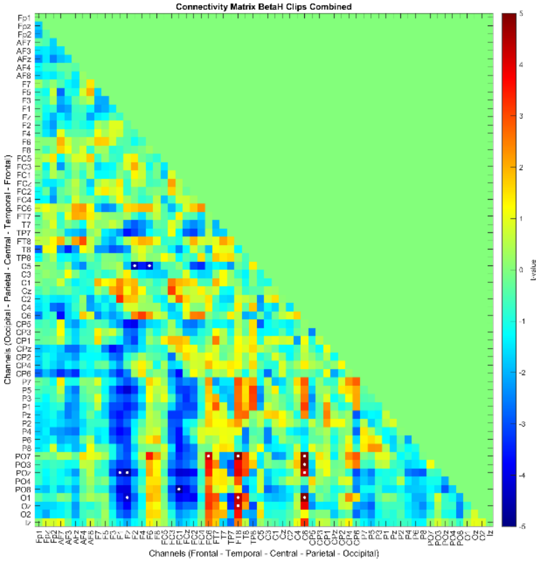


Clips Combined

Analysis was done between control and stress conditions for each clip separately and for the combined clips. The color of connections denotes the direction of connectivity alterations, where red denotes increased connectivity in the stress condition and blue reduced connectivity compared to the control condition. Significant points are highlighted with a white dot. The order of channels is Frontal - Temporal - Central - Parietal - Occipital.

### Low Gamma Range Connectivity Matrix (Clip 1 – 4; Clips Combined)

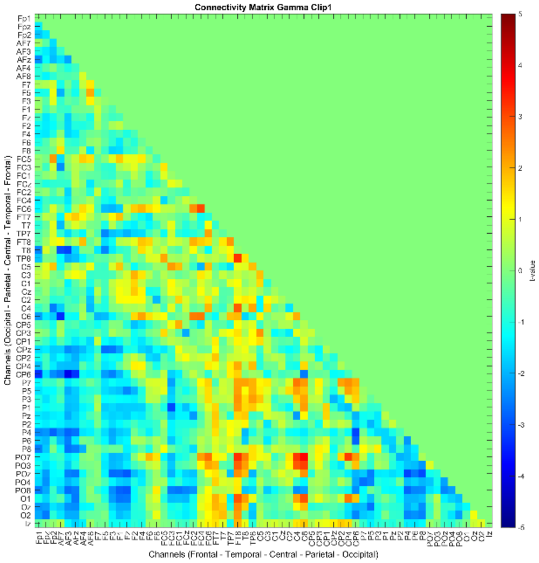


Clip 1

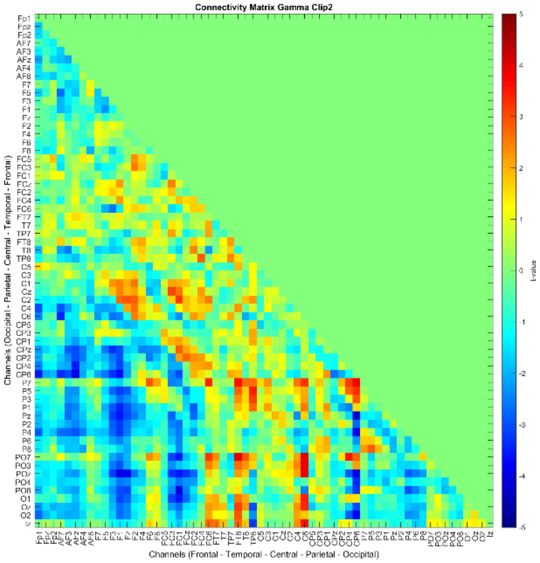


Clip 2

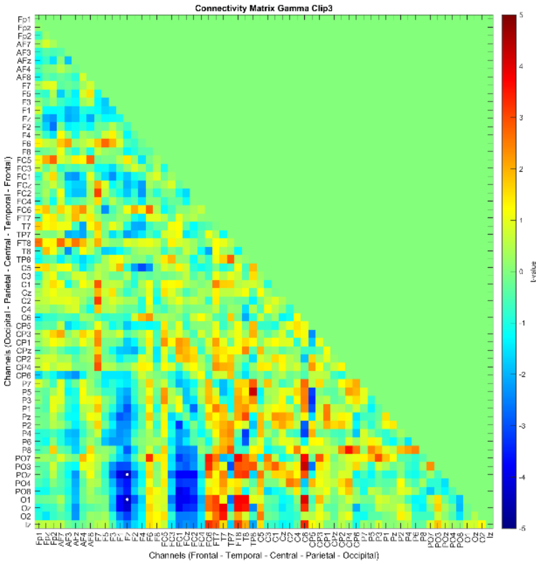


Clip 3

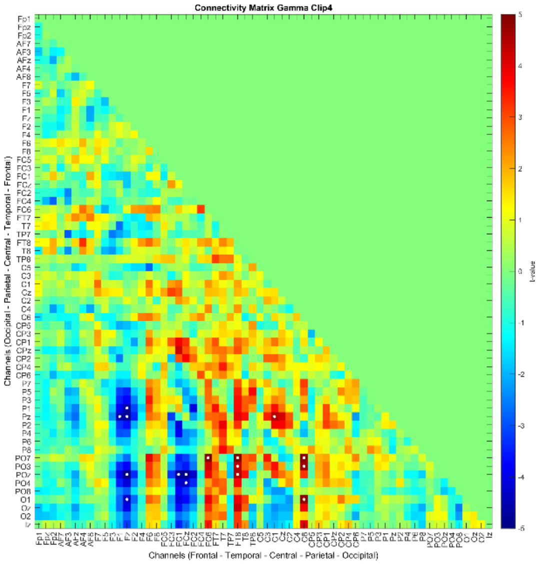


Clip 4

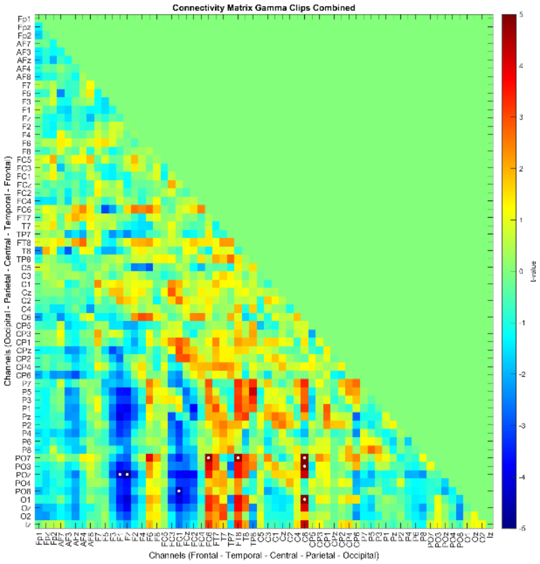


Clips Combined

Analysis was done between control and stress conditions for each clip separately and for the combined clips. The color of connections denotes the direction of connectivity alterations, where red denotes increased connectivity in the stress condition and blue reduced connectivity compared to the control condition. Significant points are highlighted with a white dot. The order of channels is Frontal - Temporal - Central - Parietal - Occipital.

### High Gamma Range Connectivity Matrix (Clip 1 – 4; Clips Combined)

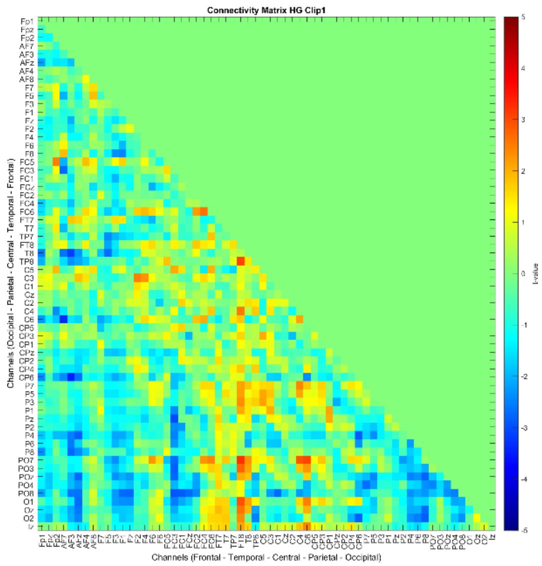


Clip 1

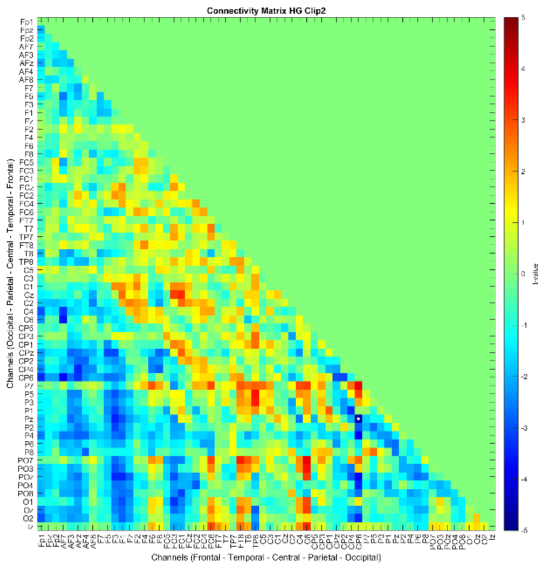


Clip 2

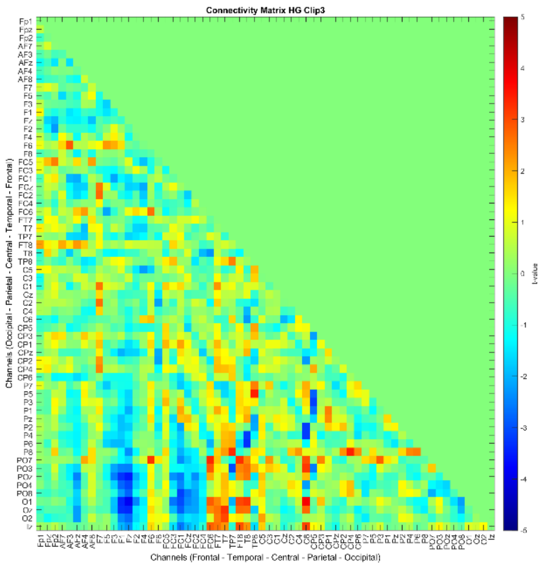


Clip 3

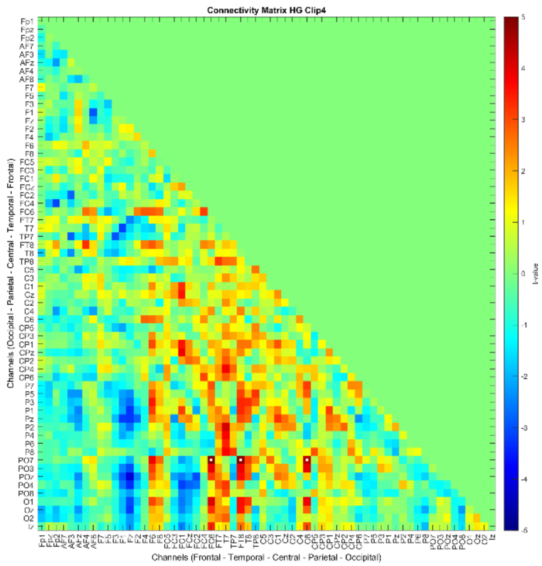


Clip 4

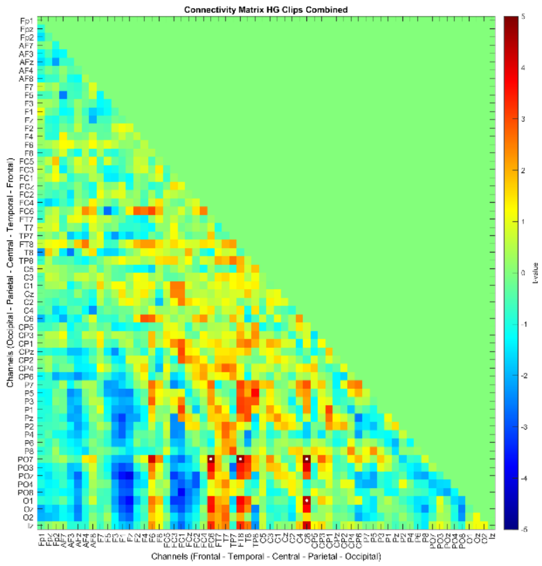


Clips Combined

Analysis was done between control and stress conditions for each clip separately and for the combined clips. The color of connections denotes the direction of connectivity alterations, where red denotes increased connectivity in the stress condition and blue reduced connectivity compared to the control condition. Significant points are highlighted with a white dot. The order of channels is Frontal - Temporal - Central - Parietal - Occipital.
