## Supplementary material for "Multimodal assessment of acute stress dynamics using an Aversive Video Paradigm (AVP)": Resting State Connectivity Matrices.pdf

### Theta Range Connectivity Matrix (Resting State – Before vs After)
### Eyes open and Eyes closed Condition

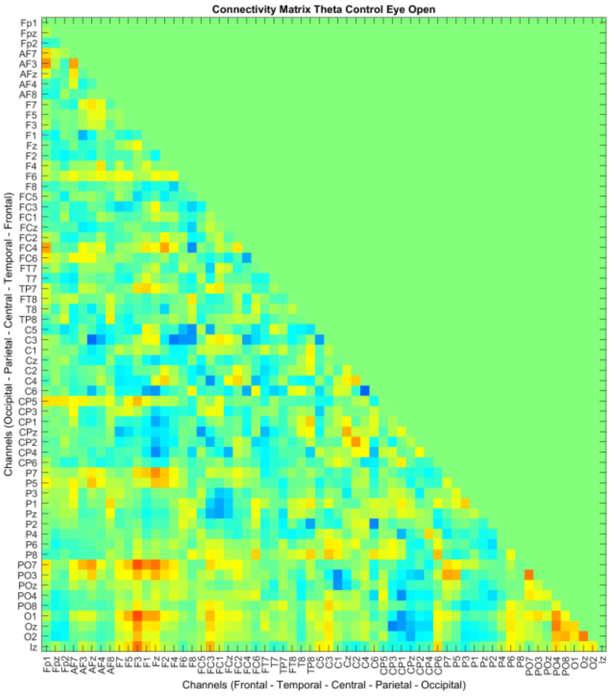


Control Eyes open

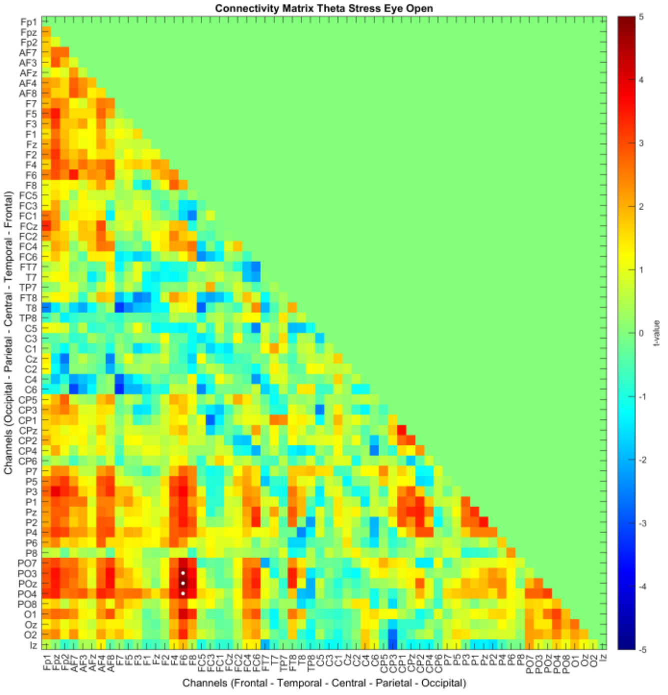


Stress Eyes open

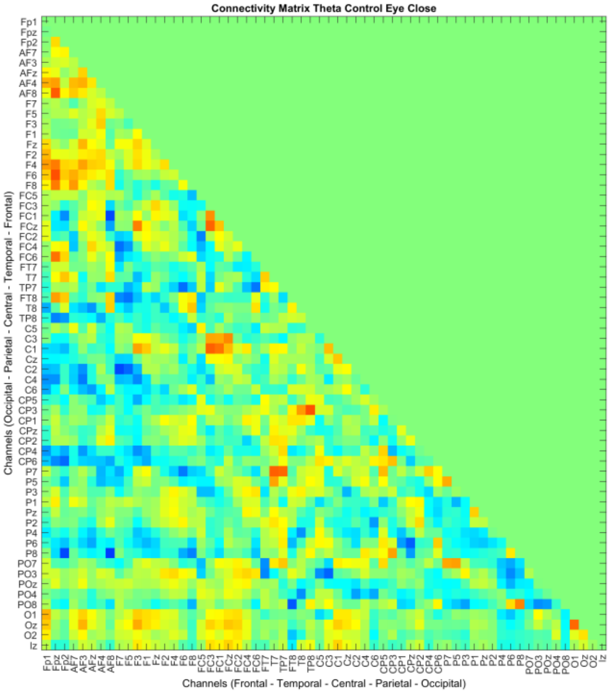


Control Eyes closed

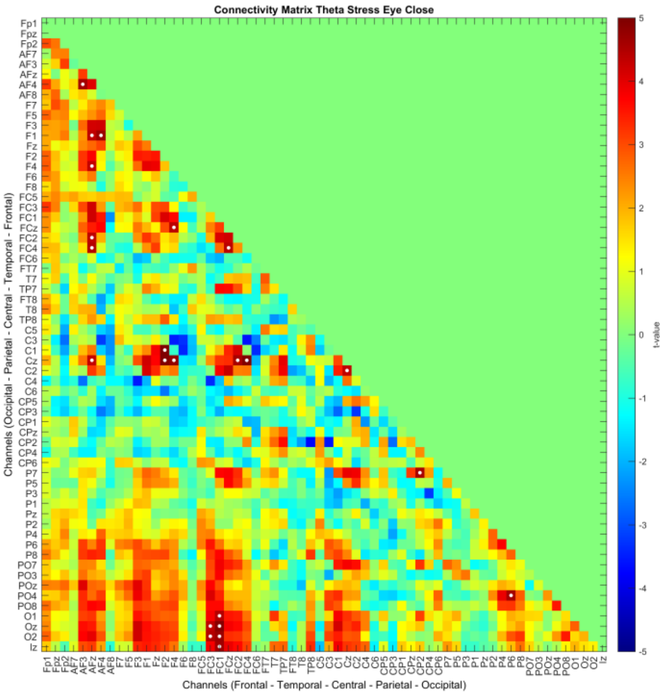


Stress Eyes closed

For Before vs After intervention analyses, every analysis was done between the before vs after intervention separately for the intervention conditions (Control, and Stress) in both, Eyes open and Eyes closed states. The color of connections denotes the direction of connectivity alterations, where red denotes increased connectivity in the resting state after compared to before intervention, and blue indicates reduced connectivity. Significant points are highlighted by a white dot. The order of channels is Frontal - Temporal - Central - Parietal - Occipital.

### Theta range connectivity matrix (Resting State – Control vs Stress) Eyes open and Eyes closed Condition

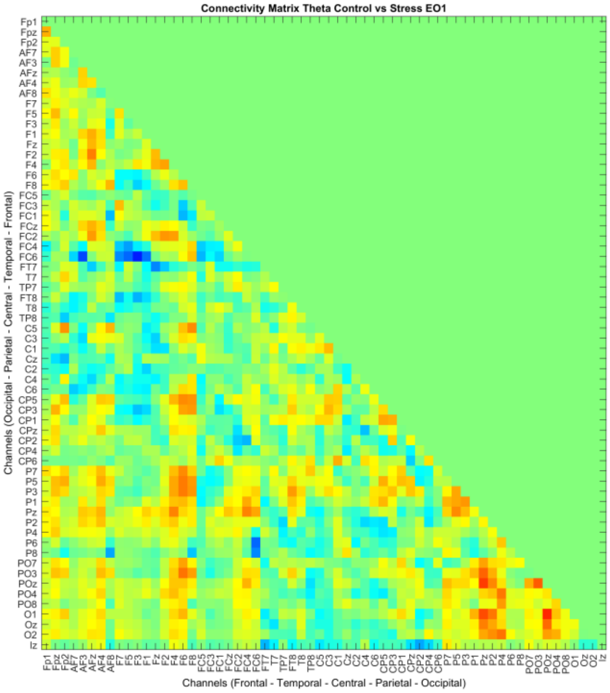


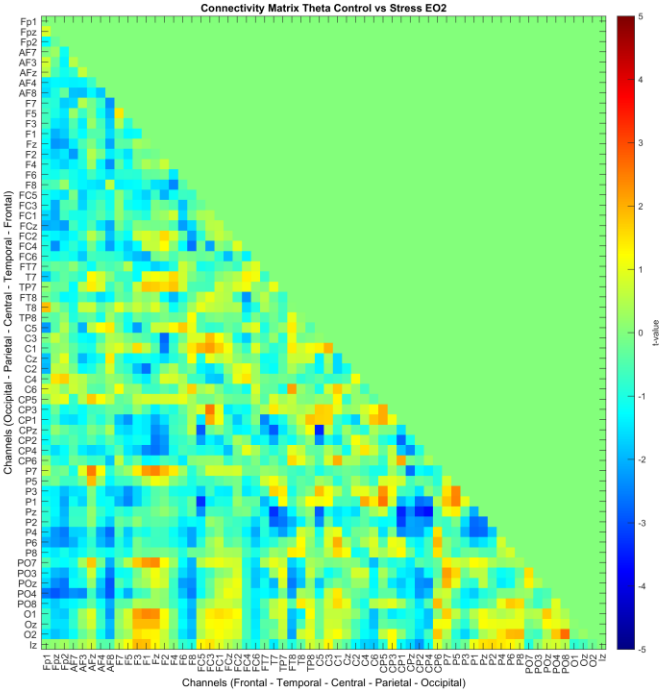


Resting State 1 - Eyes open

Resting State 2 - Eyes open

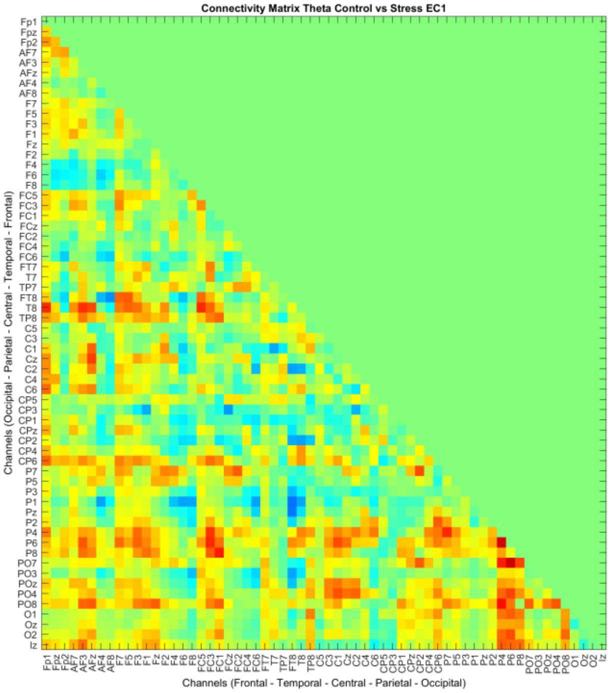


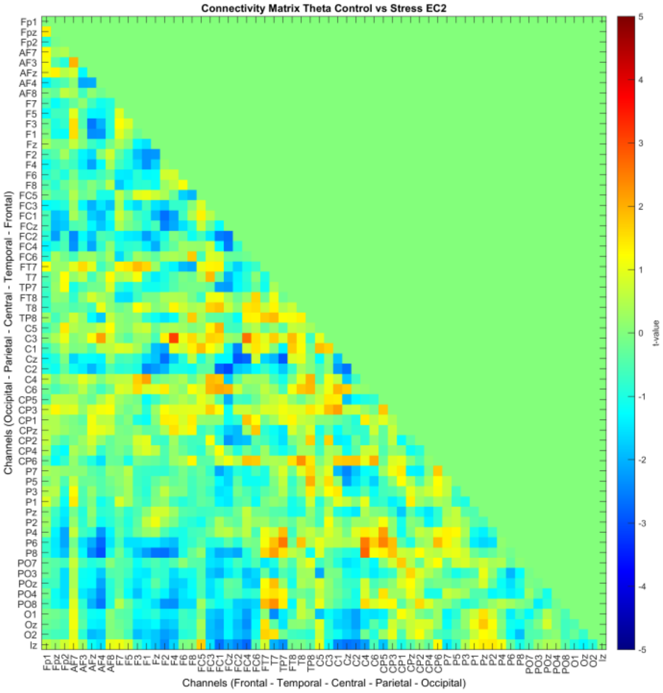


Resting State 1 - Eyes closed

Resting State 2 - Eyes closed

For Control vs Stress intervention analyses, every analysis was done between the Control vs Stress conditions for each resting state (before and after intervention) separately in Eyes Open (EO) and Eyes Closed (EC) states. The color of connections denotes the direction of connectivity alterations, where red denotes increased connectivity in the stress condition, and blue indicates reduced connectivity in stress conditions as compared to control. No significant points were found in this analysis. The order of channels is similar to the above.

### Alpha range connectivity matrix (Resting State – Before vs After)
### Eyes open and Eyes closed Condition

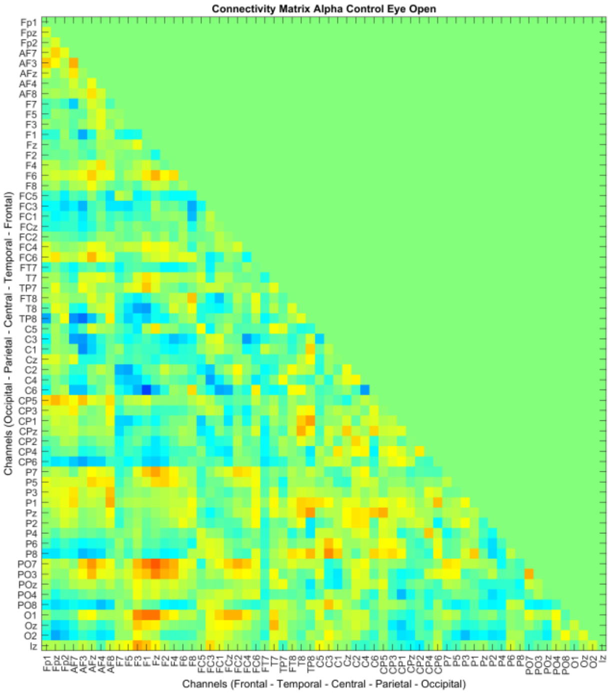


Control Eyes open

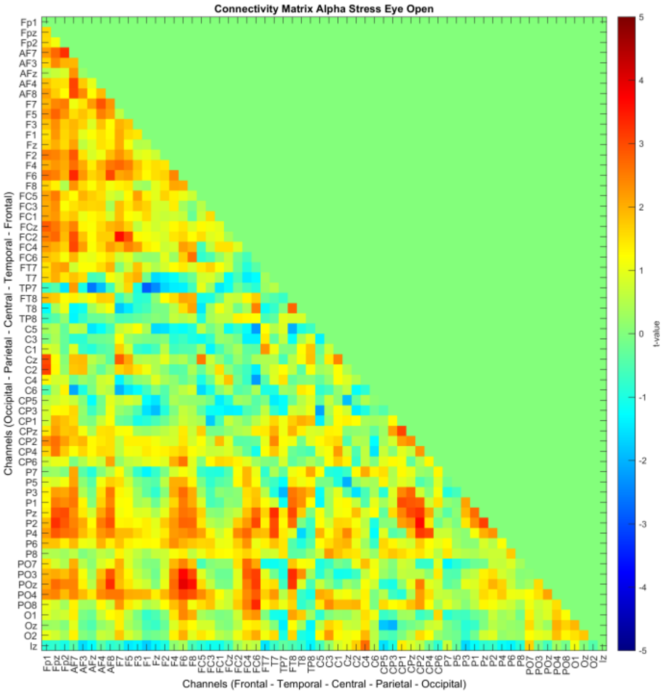


Stress Eyes open

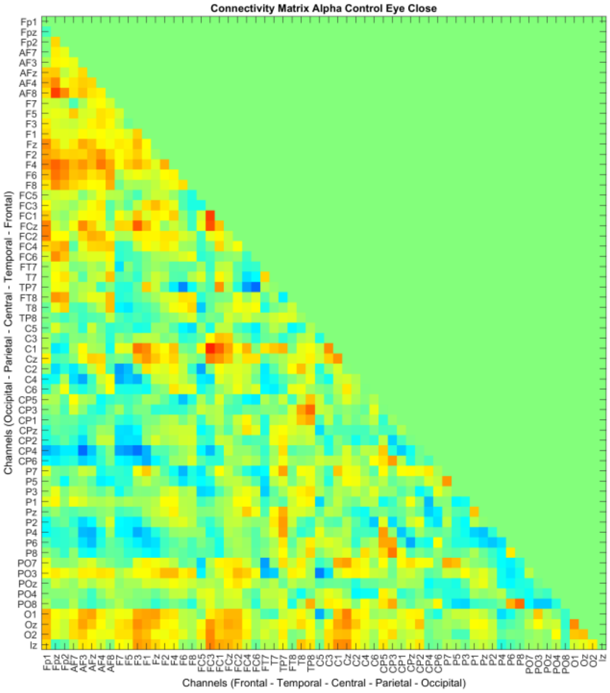


Control Eyes closed

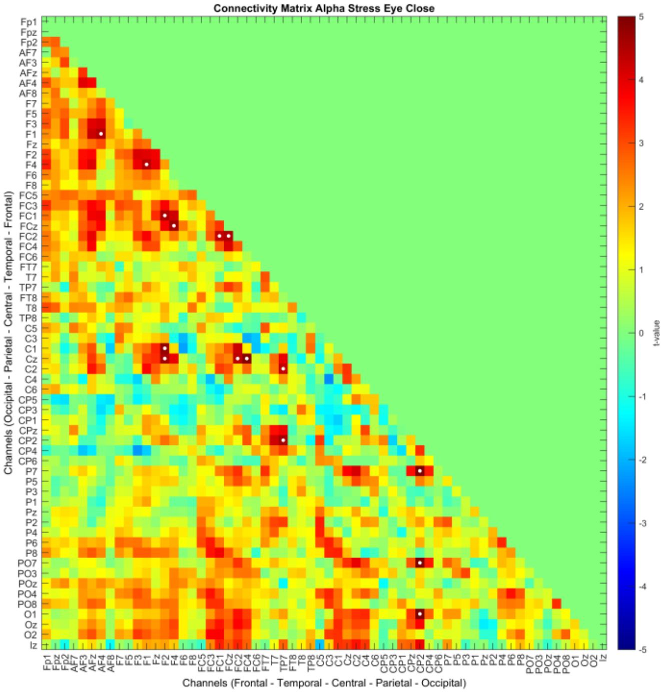


Stress Eyes closed

For Before vs After intervention analyses, every analysis was done between the before vs after intervention separately for the intervention conditions (Control, and Stress) in both, Eyes open and Eyes closed states. The color of connections denotes the direction of connectivity alterations, where red denotes increased connectivity in the resting state after compared to before intervention, and blue indicates reduced connectivity. Significant points are highlighted by a white dot. The order of channels is Frontal - Temporal - Central - Parietal - Occipital.

### Alpha range connectivity matrix (Resting State – Control vs Stress) Eyes open and Eyes closed Condition

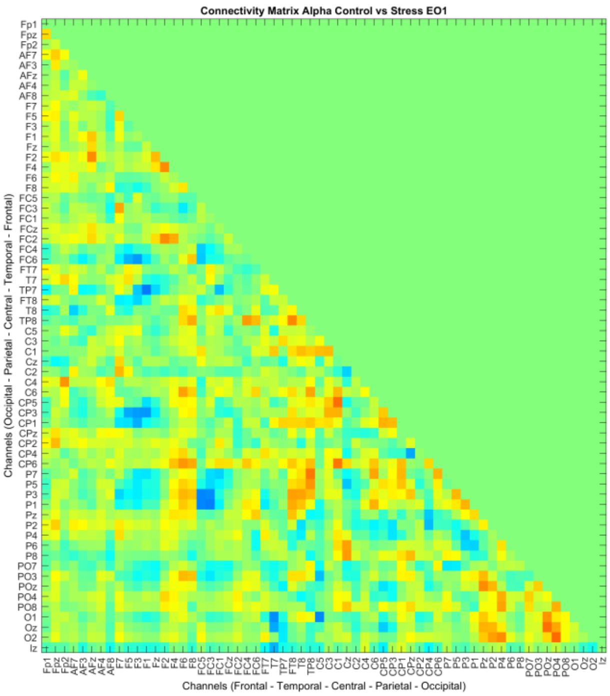


Resting State 1 - Eyes open

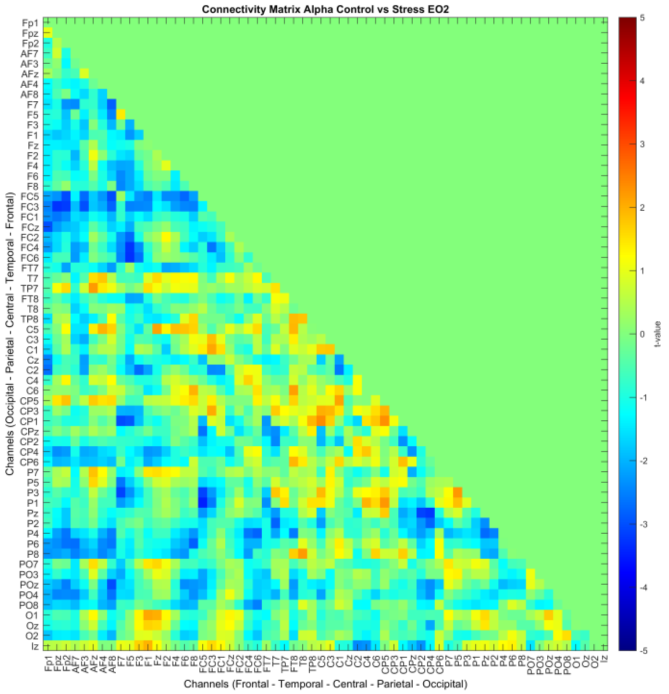


Resting State 2 - Eyes open

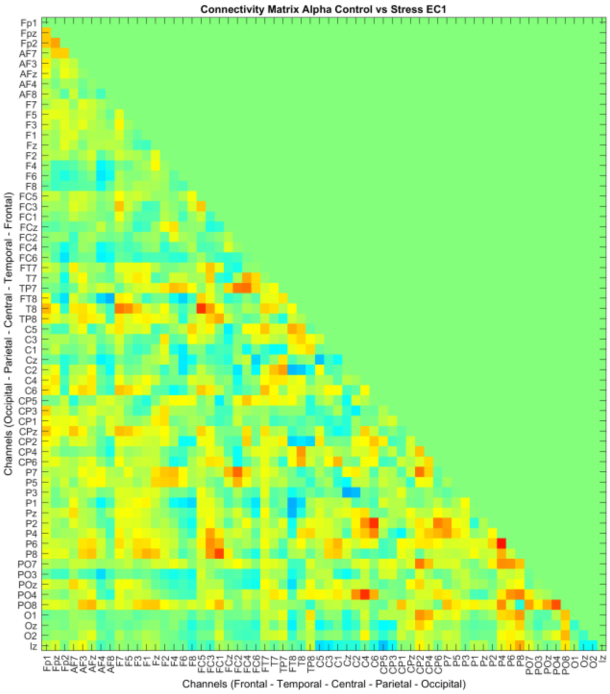


Resting State 1 - Eyes closed

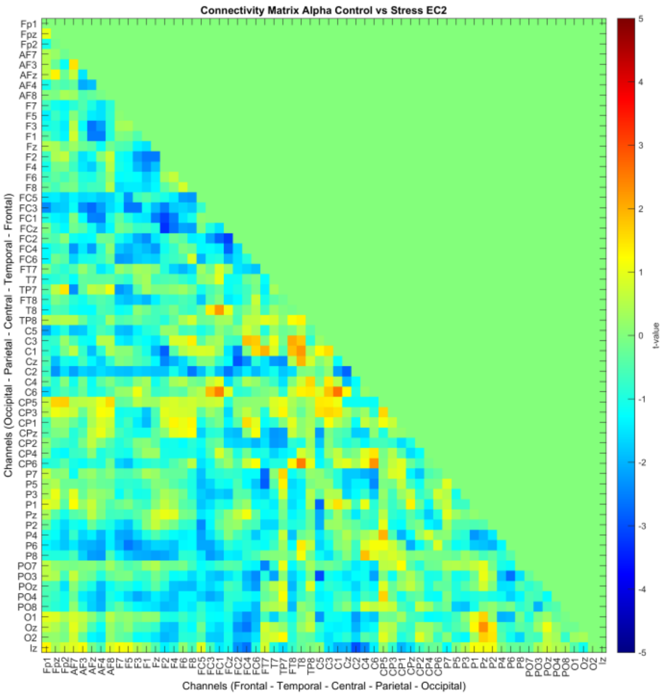


Resting State 2 - Eyes closed

For Control vs Stress intervention analyses, every analysis was done between the Control vs Stress conditions for each resting state (before and after intervention) separately in Eyes Open (EO) and Eyes Closed (EC) states. The color of connections denotes the direction of connectivity alterations, where red denotes increased connectivity in the stress condition, and blue indicates reduced connectivity in stress conditions as compared to control. No significant points were found in this analysis. The order of channels is similar to the above.

### Low Beta range connectivity matrix (Resting State – Before vs After) Eyes open and Eyes closed Condition

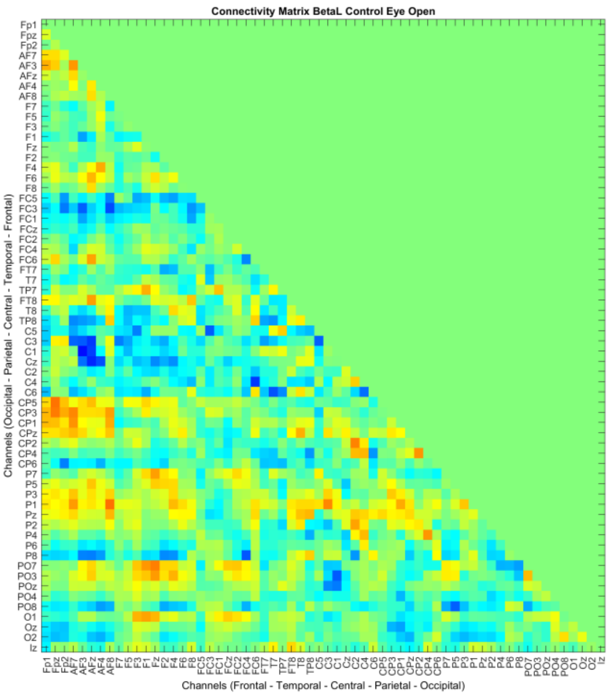


Control Eyes open

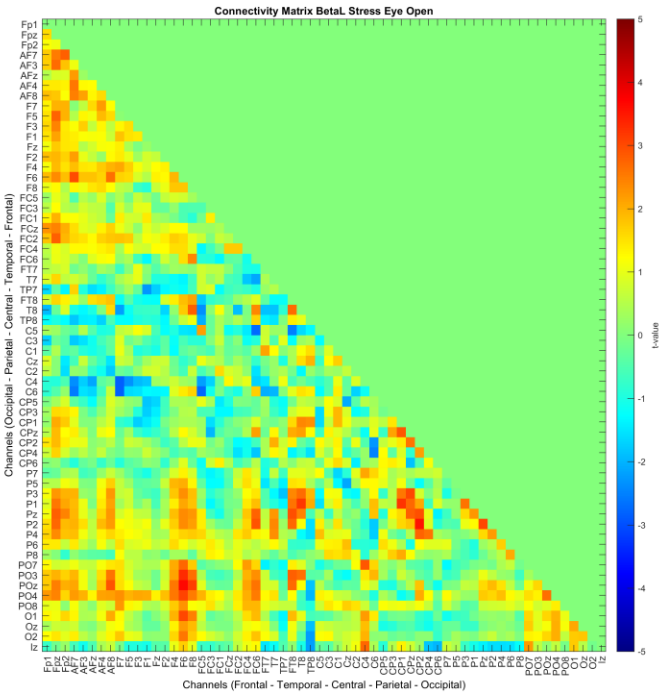


Stress Eyes open

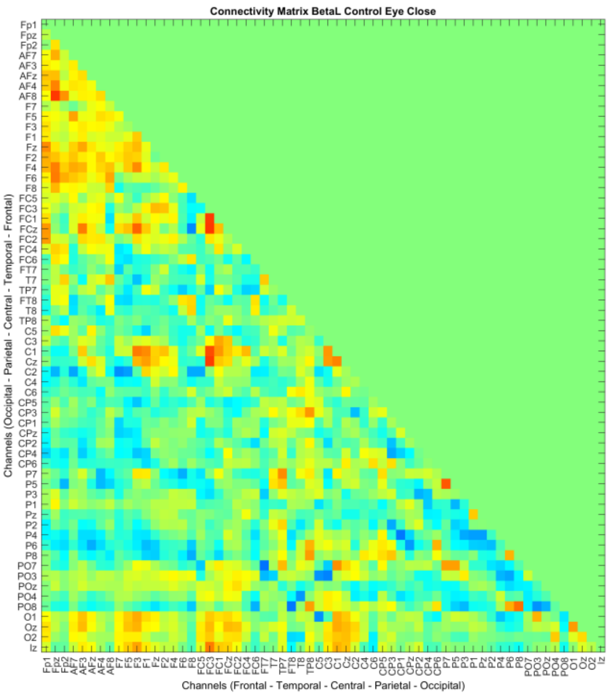


Control Eyes closed

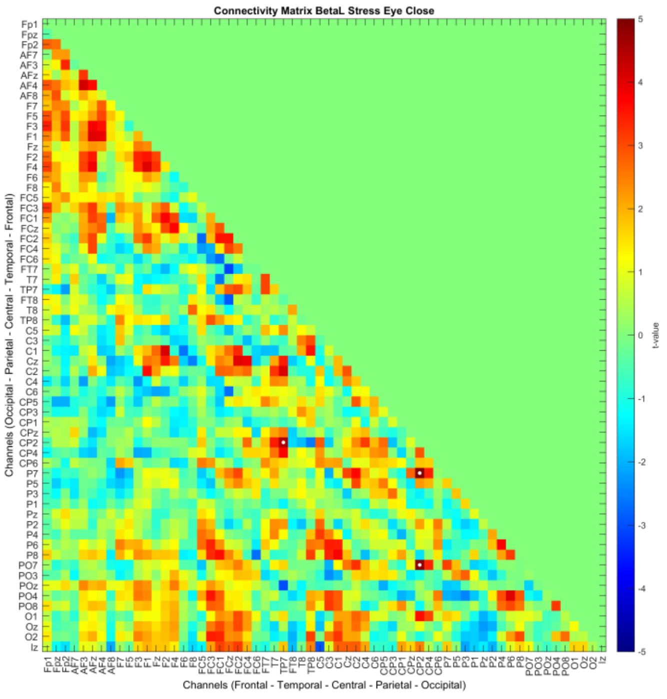


Stress Eyes closed

For Before vs After intervention analyses, every analysis was done between the before vs after intervention separately for the intervention conditions (Control, and Stress) in both, Eyes open and Eyes closed states. The color of connections denotes the direction of connectivity alterations, where red denotes increased connectivity in the resting state after compared to before intervention, and blue indicates reduced connectivity. Significant points are highlighted by a white dot. The order of channels is Frontal - Temporal - Central - Parietal - Occipital.

### Low Beta range connectivity matrix (Resting State – Control vs Stress) Eyes open and Eyes closed Condition

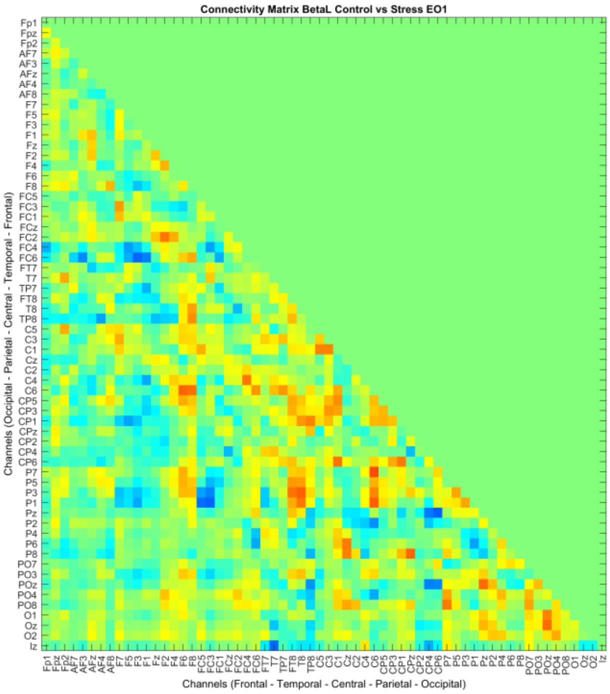


Resting State 1 - Eyes open

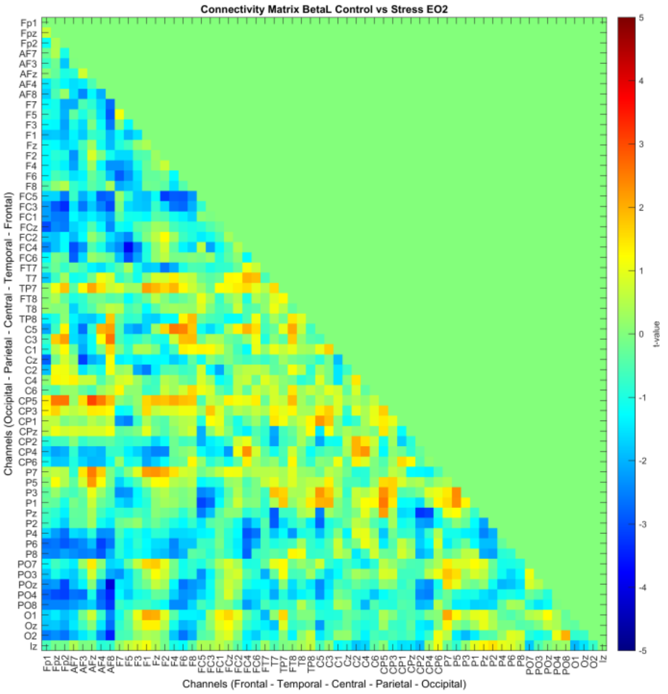


Resting State 2 - Eyes open

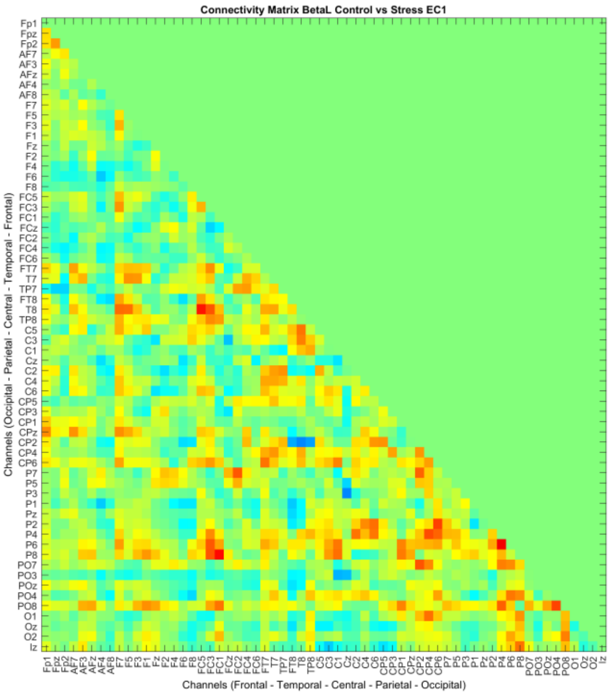


Resting State 1 - Eyes closed

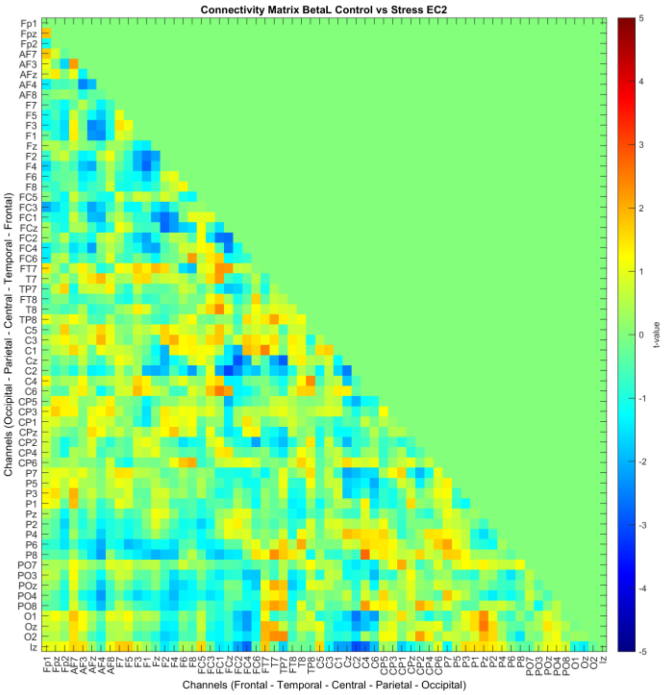


Resting State 2 - Eyes closed

For Control vs Stress intervention analyses, every analysis was done between the Control vs Stress conditions for each resting state (before and after intervention) separately in Eyes Open (EO) and Eyes Closed (EC) states. The color of connections denotes the direction of connectivity alterations, where red denotes increased connectivity in the stress condition, and blue indicates reduced connectivity in stress conditions as compared to control. No significant points were found in this analysis. The order of channels is similar to the above.

### High Beta range connectivity matrix (Resting State – Before vs After) Eyes open and Eyes closed Condition

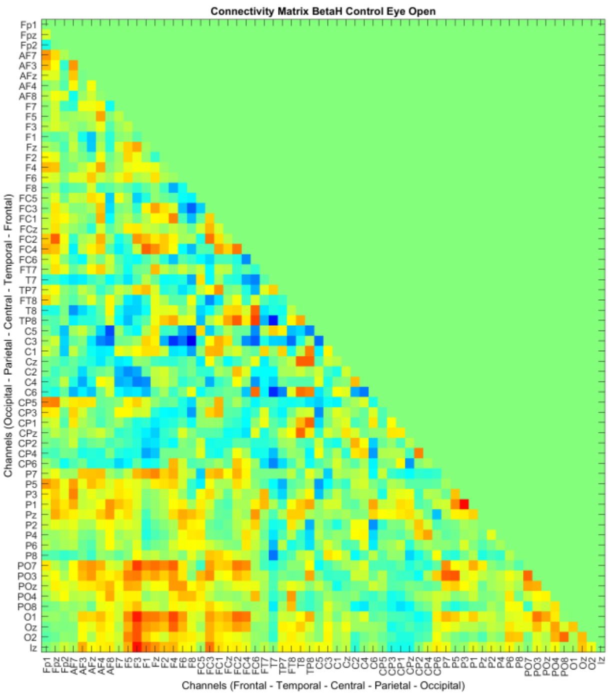


Control Eyes open

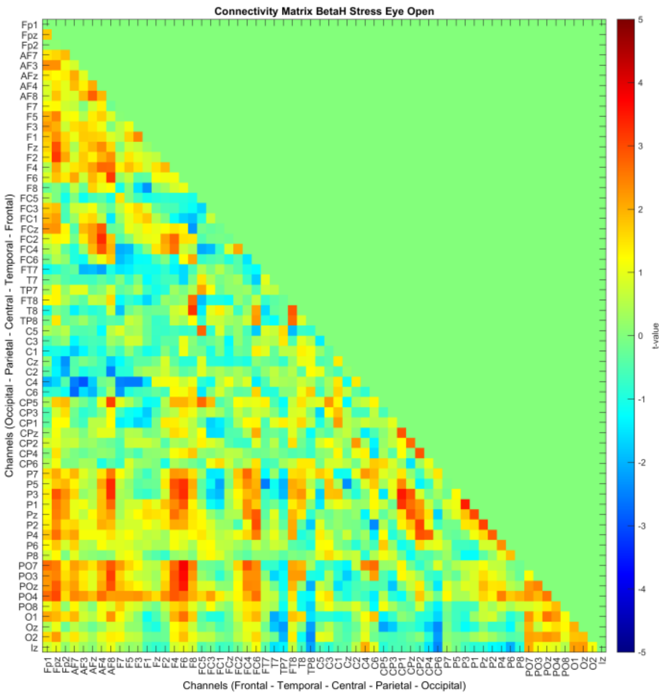


Stress Eyes open

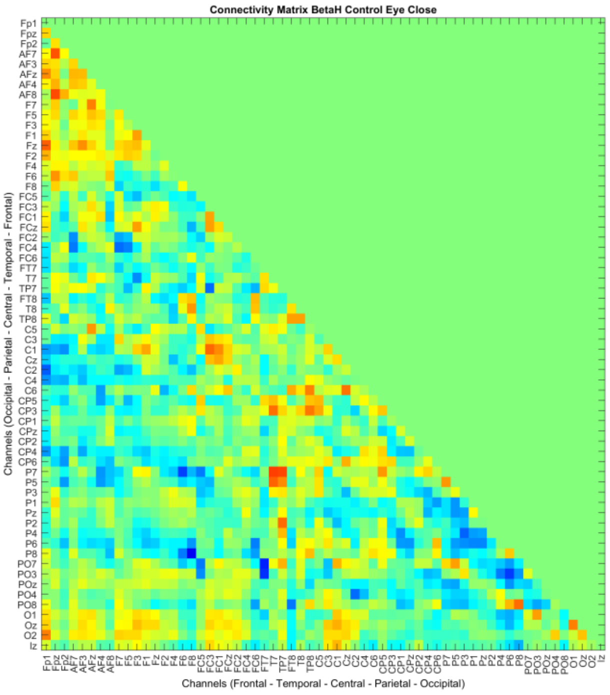


Control Eyes closed

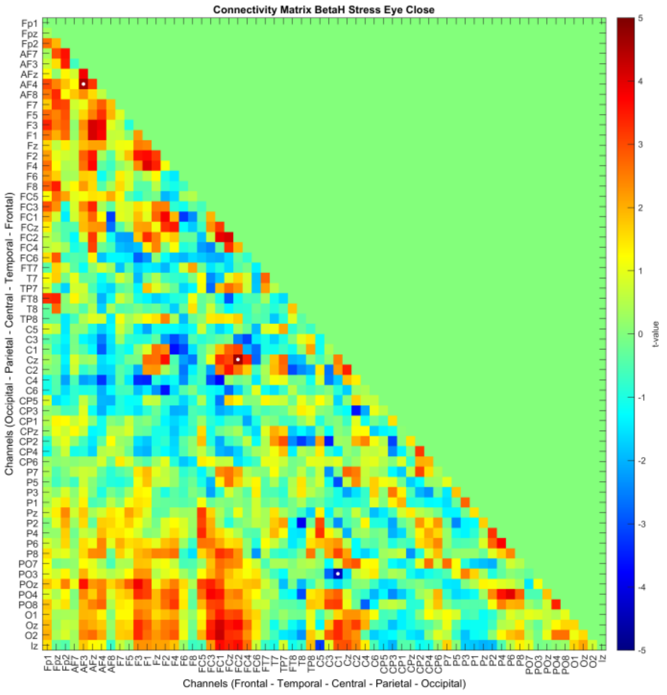


Stress Eyes closed

For Before vs After intervention analyses, every analysis was done between the before vs after intervention separately for the intervention conditions (Control, and Stress) in both, Eyes open and Eyes closed states. The color of connections denotes the direction of connectivity alterations, where red denotes increased connectivity in the resting state after compared to before intervention, and blue indicates reduced connectivity. Significant points are highlighted by a white dot. The order of channels is Frontal - Temporal - Central - Parietal - Occipital.

### High Beta range connectivity matrix (Resting State – Control vs Stress) Eyes open and Eyes closed Condition

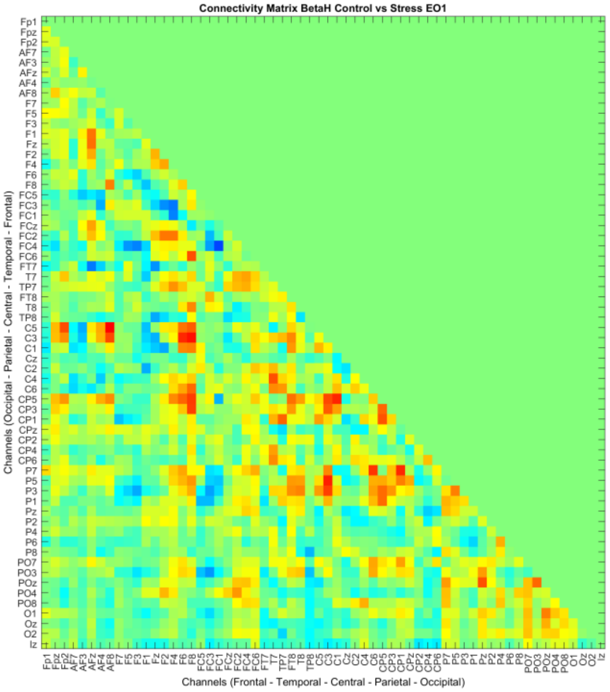


Resting State 1 - Eyes open

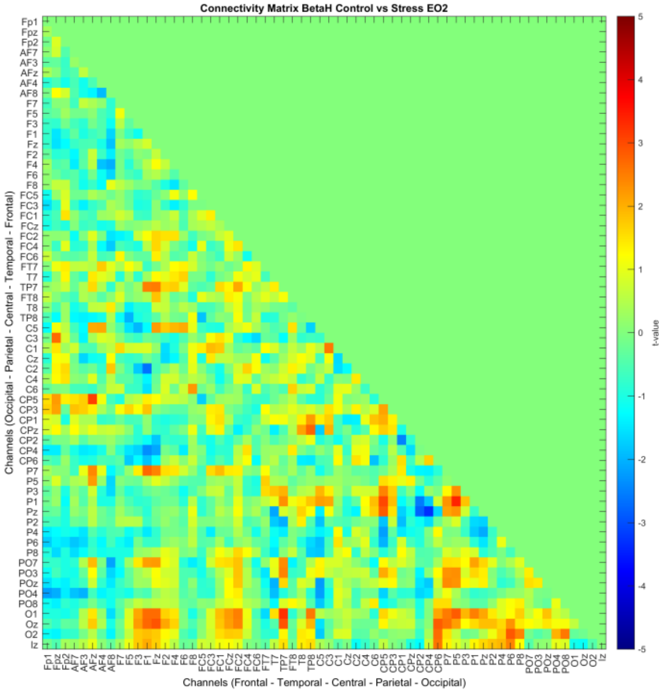


Resting State 2 - Eyes open

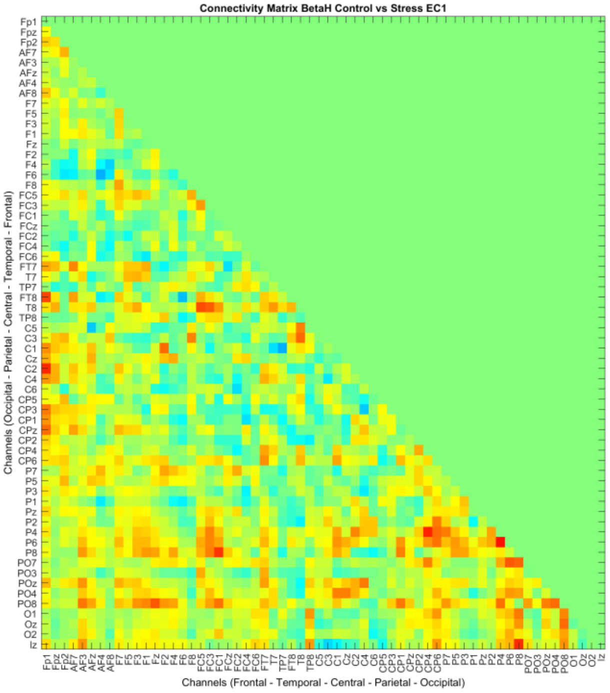


Resting State 1 - Eyes closed

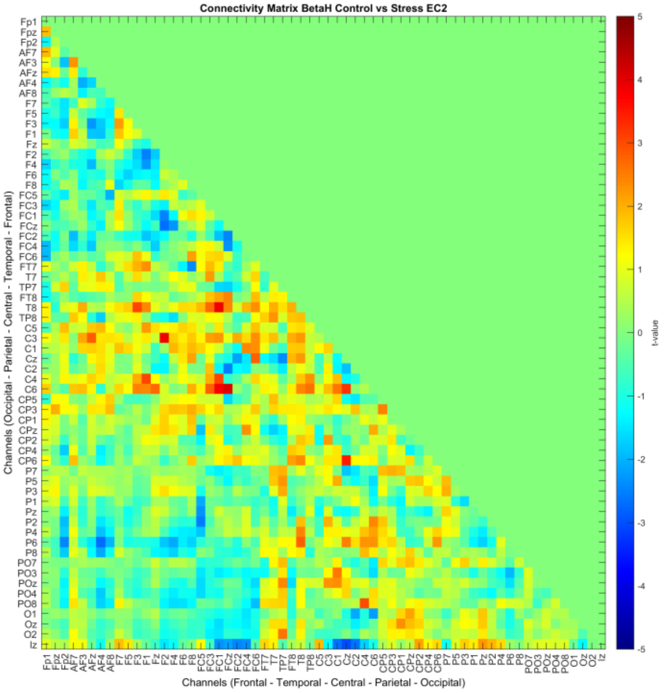


Resting State 2 - Eyes closed

For Control vs Stress intervention analyses, every analysis was done between the Control vs Stress conditions for each resting state (before and after intervention) separately in Eyes Open (EO) and Eyes Closed (EC) states. The color of connections denotes the direction of connectivity alterations, where red denotes increased connectivity in the stress condition, and blue indicates reduced connectivity in stress conditions as compared to control. No significant points were found in this analysis. The order of channels is similar to the above.

### Low Gamma range connectivity matrix (Resting State – Before vs After) Eyes open and Eyes closed Condition

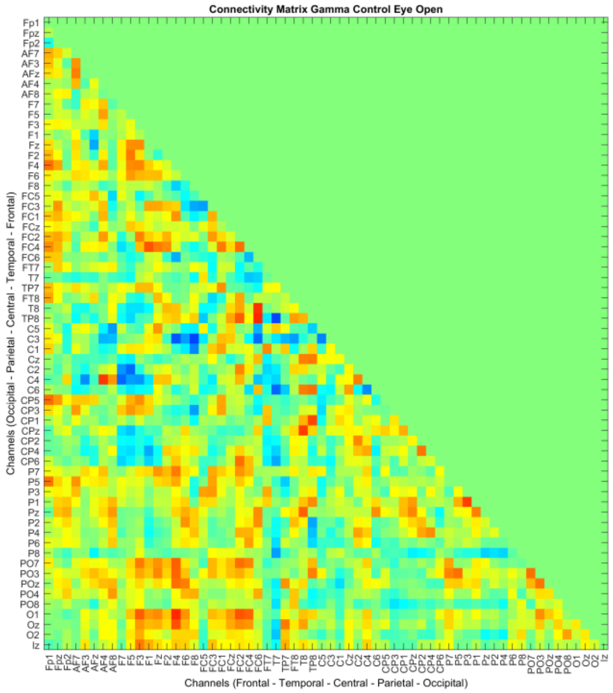


Control Eyes open

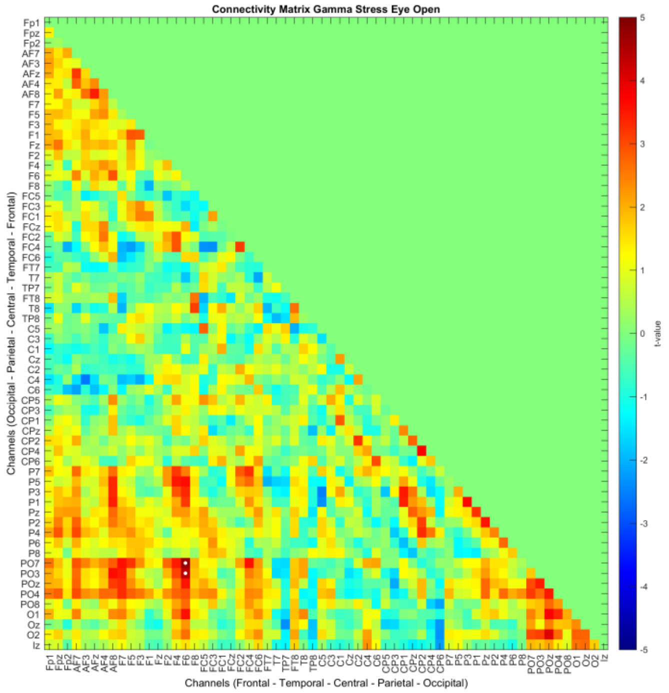


Stress Eyes open

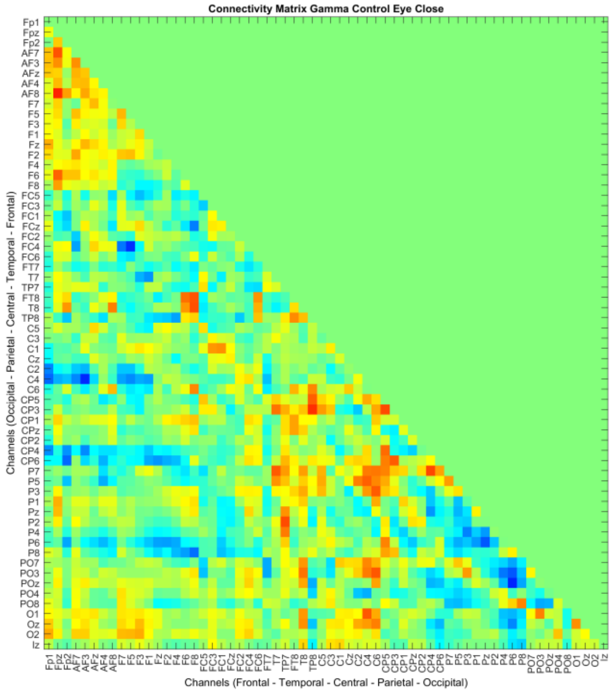


Control Eyes closed

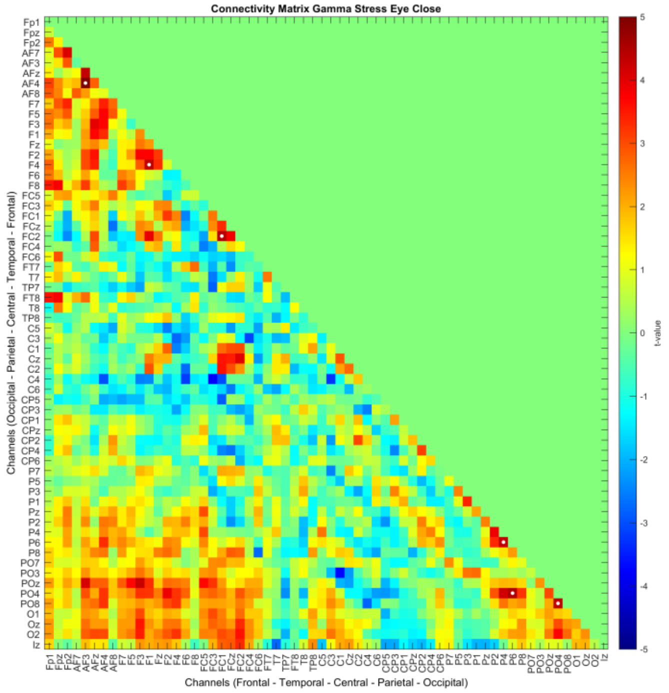


Stress Eyes closed

For Before vs After intervention analyses, every analysis was done between the before vs after intervention separately for the intervention conditions (Control, and Stress) in both, Eyes open and Eyes closed states. The color of connections denotes the direction of connectivity alterations, where red denotes increased connectivity in the resting state after compared to before intervention, and blue indicates reduced connectivity. Significant points are highlighted by a white dot. The order of channels is Frontal - Temporal - Central - Parietal - Occipital.

### Low Gamma range connectivity matrix (Resting State – Control vs Stress) Eyes open and Eyes closed Condition

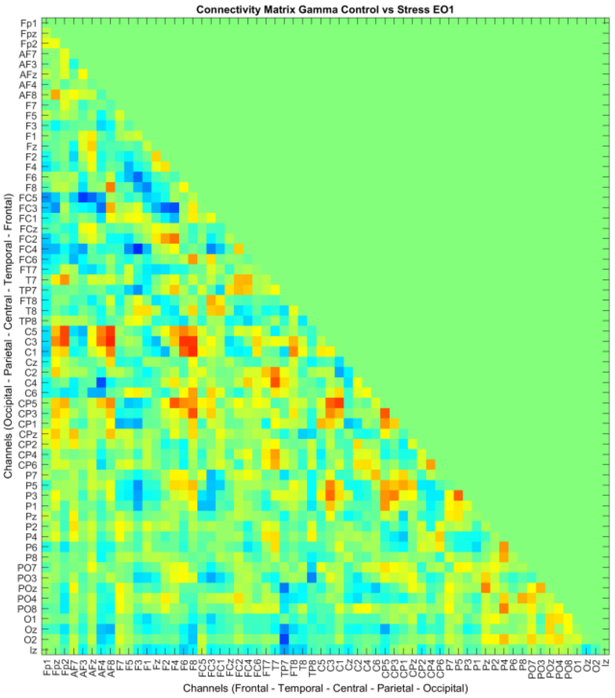


Resting State 1 - Eyes open

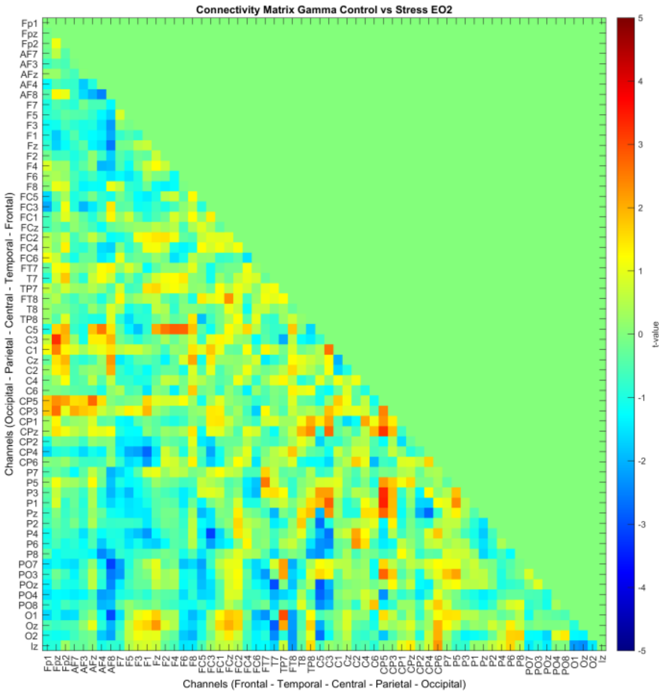


Resting State 2 - Eyes open

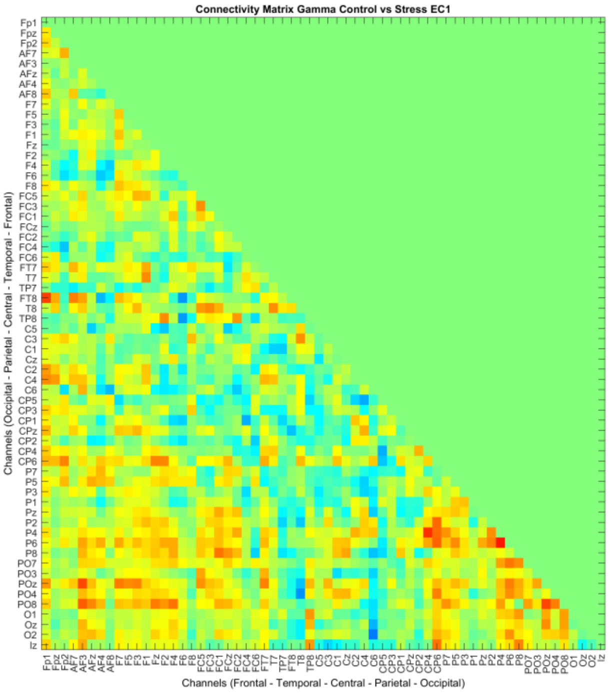


Resting State 1 - Eyes closed

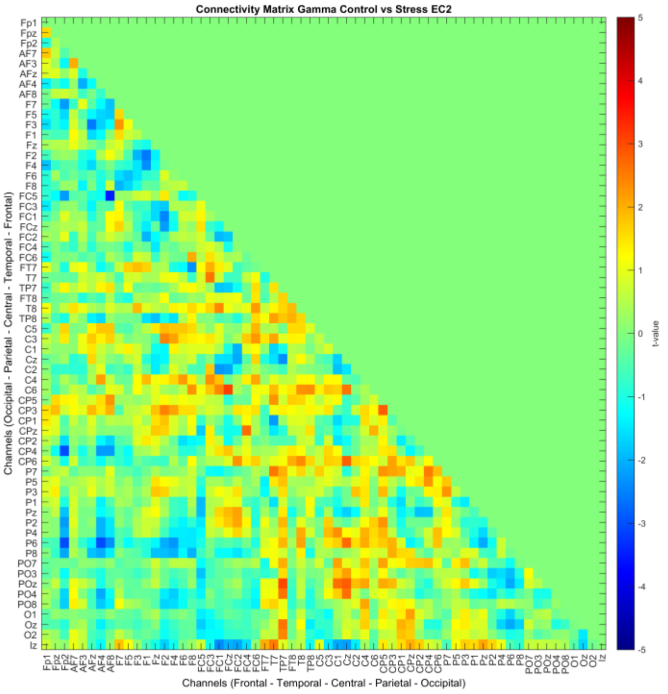


Resting State 2 - Eyes closed

For Control vs Stress intervention analyses, every analysis was done between the Control vs Stress conditions for each resting state (before and after intervention) separately in Eyes Open (EO) and Eyes Closed (EC) states. The color of connections denotes the direction of connectivity alterations, where red denotes increased connectivity in the stress condition, and blue indicates reduced connectivity in stress conditions as compared to control. No significant points were found in this analysis. The order of channels is similar to the above.

### High Gamma range connectivity matrix (Resting State – Before vs After) Eyes open and Eyes closed Condition

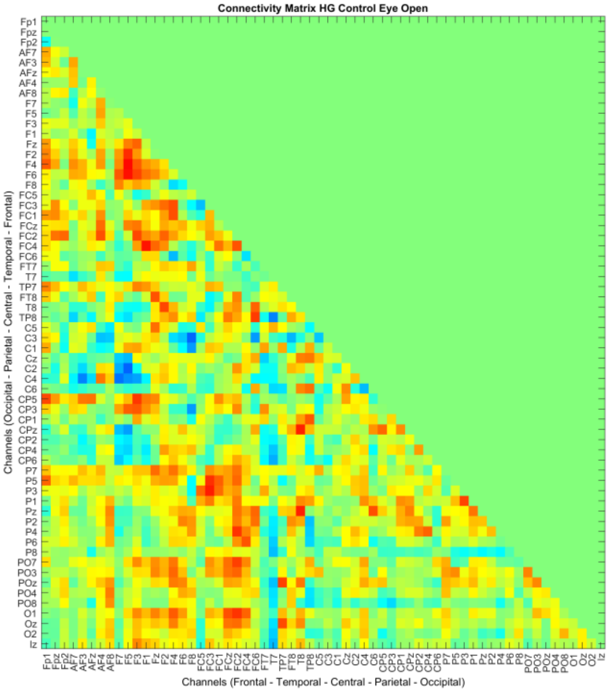


Control Eyes open

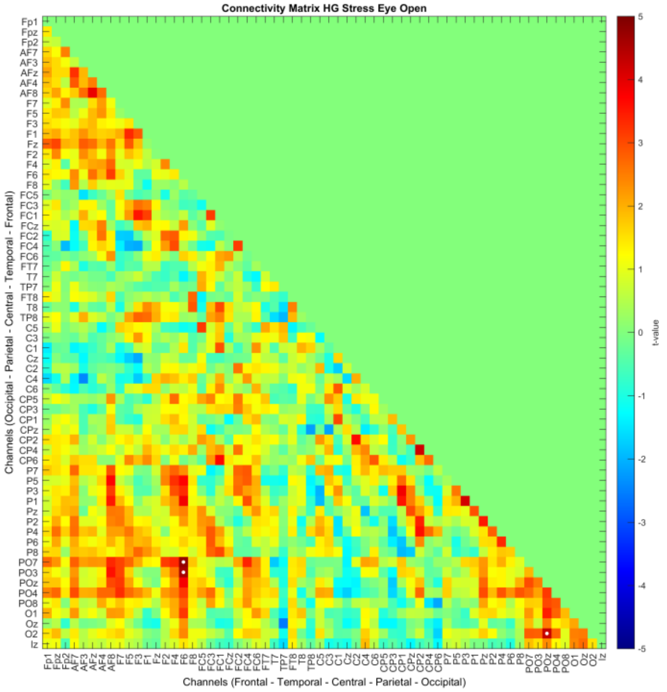


Stress Eyes open

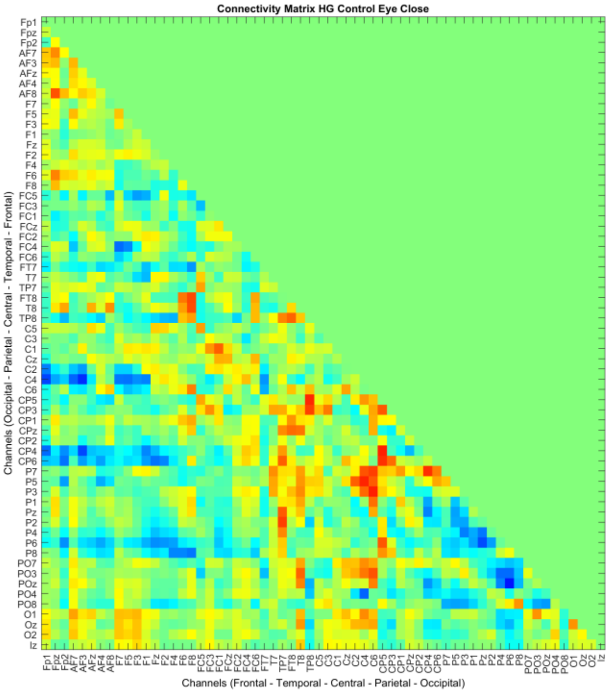


Control Eyes closed

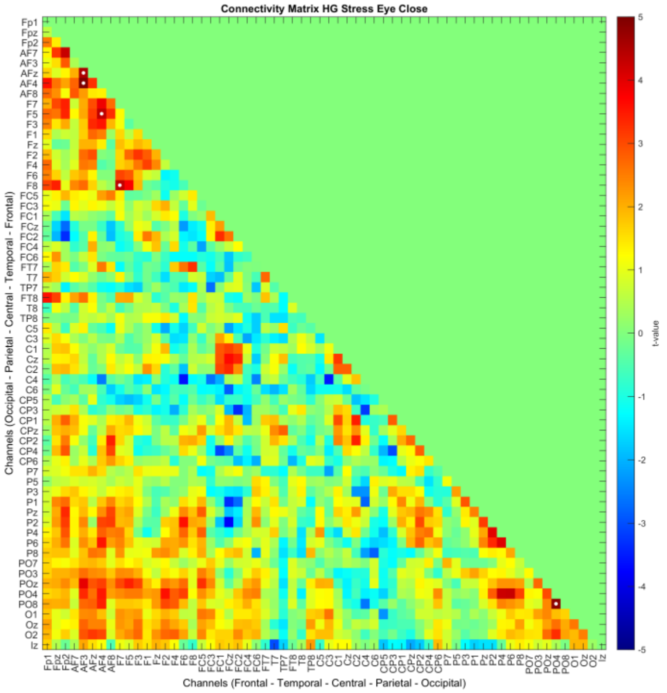


Stress Eyes closed

For Before vs After intervention analyses, every analysis was done between the before vs after intervention separately for the intervention conditions (Control, and Stress) in both, Eyes open and Eyes closed states. The color of connections denotes the direction of connectivity alterations, where red denotes increased connectivity in the resting state after compared to before intervention, and blue indicates reduced connectivity. Significant points are highlighted by a white dot. The order of channels is Frontal - Temporal - Central - Parietal - Occipital.

### High Gamma range connectivity matrix (Resting State – Control vs Stress) Eyes open and Eyes closed Condition

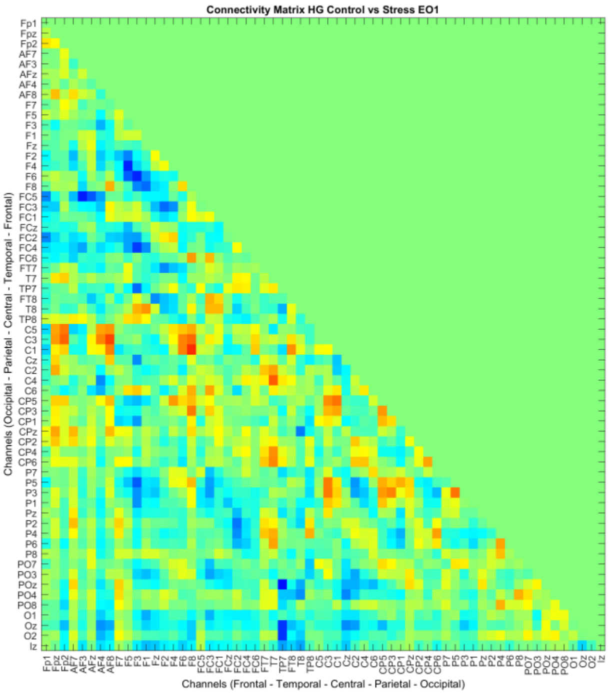


Resting State 1 - Eyes open

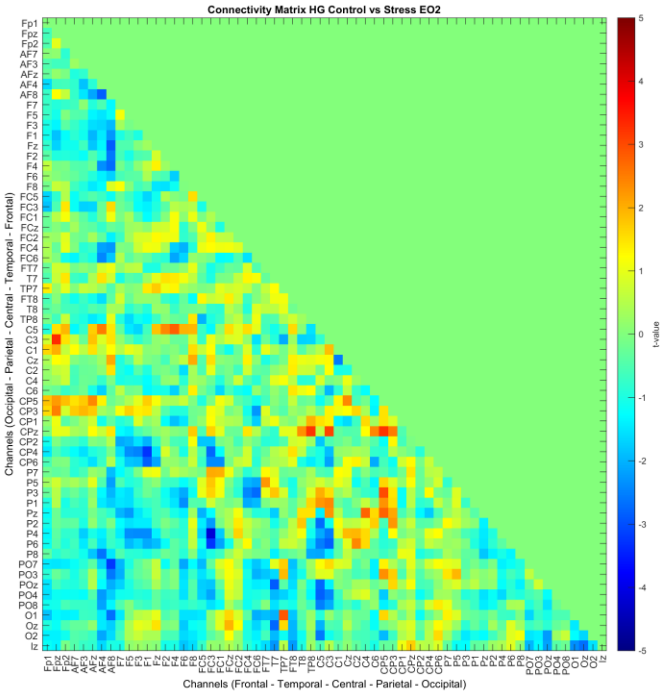


Resting State 2 - Eyes open

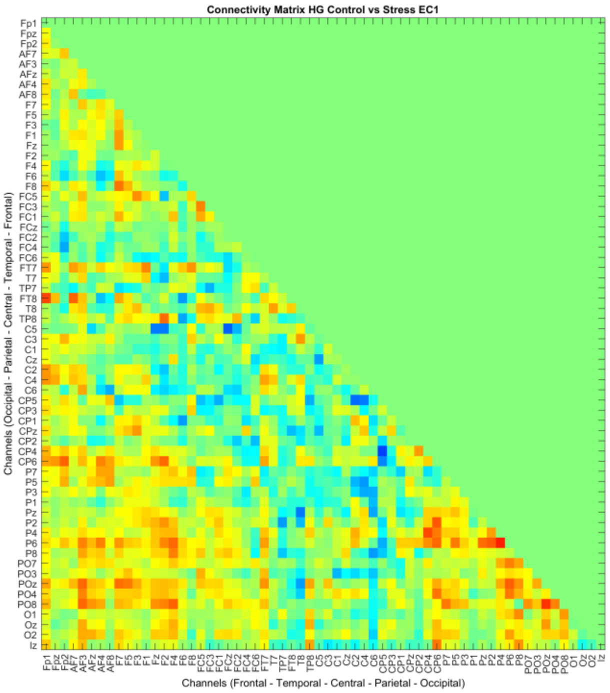


Resting State 1 - Eyes closed

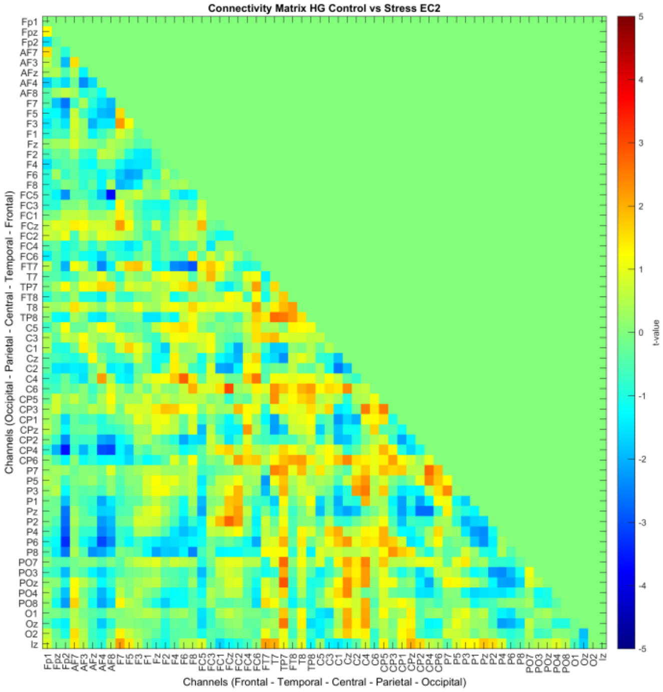


Resting State 2 - Eyes closed

For Control vs Stress intervention analyses, every analysis was done between the Control vs Stress conditions for each resting state (before and after intervention) separately in Eyes Open (EO) and Eyes Closed (EC) states. The color of connections denotes the direction of connectivity alterations, where red denotes increased connectivity in the stress condition, and blue indicates reduced connectivity in stress conditions as compared to control. No significant points were found in this analysis. The order of channels is similar to the above.
