## Supplementary material for "Multimodal assessment of acute stress dynamics using an Aversive Video Paradigm (AVP)": Supplementary Sheet.pdf

*Table 1 Cluster information of the EEG power analysis during movie clip presentation. Statistics were performed between control and stress conditions for each clip individually and for all clips combined. Cluster-based permutation testing was employed to identify significant clusters in the data. This table shows information about significant clusters found in each analysis in the frequency band of interest. t-values indicate the sum of the t-values of the whole cluster, the p-value indicates the significance level, and the size of the cluster is shown by the number of the involved electrodes. The direction of the effect is indicated by the sign of the t-value, where a positive t-value indicates increased power in stress conditions and a negative t-value indicates decreased power in stress relative to control conditions. Comparisons which would include absent clusters or clusters below the threshold are not shown.*

| | | ***All clips combined*** | ***Clip 1*** | ***Clip 2*** | ***Clip 3*** | ***Clip4*** |
|---|---|---|---|---|---|---|
| **Theta (4-8 Hz)** | P value | <0.001 | 0.0034 | 0.0014 | 0.0039 | <.001 |
| | t-value | -101.07 | -53.92 | -72.69 | -36.67 | -140.15 |
| | No. of electrodes | 32 | 19 | 26 | 13 | 40 |
| **Alpha (8-13 Hz)** | P value | <0.001 | <0.001 | <0.001 | <0.001 | <.001 |
| | t-value | -287.02 | -280.82 | -267.84 | -165.71 | -301 |
| | No. of electrode | 61 | 62 | 61 | 52 | 60 |
| **Low Beta (13-15 Hz )** | P value | <0.001 | <0.001 | 0.0016 | 0.0017 | <.001 |
| | t-value | -113.87 | -159.22 | -70.99 | -49.39 | -99.36 |
| | No. of electrode | 34 | 47 | 21 | 18 | 31 |
| **High Beta (23-32 Hz )** | No significant clusters were observed in this frequency range | | | | | |
| **Low Gamma (32-50 Hz)** | P value | - | - | 0.002 | - | <.001 |
| | t-value | - | - | 71.52 | - | 94.27 |
| | No. of electrode | - | - | 26 | - | 33 |
| **High Gamma (50-80 Hz)** | P value | - | - | - | - | 0.0019 |
| | t-value | - | - | - | - | 100.34 |
| | No. of electrode | - | - | - | - | 33 |

*Table 2 Cluster information for EEG power analysis during resting states comparing before and after intervention. Statistics were done between RS2 and RS1 for the stress condition in EO and EC states. Cluster-based permutation testing was employed to identify significant clusters in the data. This table shows cluster information for significant clusters found in each analysis in the frequency band of interest. t-values indicate the combined t-values of the whole cluster, the p-value indicates the significance level, and the size of the cluster is shown by the number of the involved electrodes. The direction of the effect is shown by the sign of the t-value, where a positive t-value indicates increased power in the after intervention state, and a negative t-value indicates decreased power in the after intervention state relative to before intervention. Comparisons which would include absent clusters or clusters below the threshold are not shown. No clusters were observed in the control condition.*

| | | ***Stress Eyes Closed*** | ***Stress Eyes Open*** |
|---|---|---|---|
| **Theta (4-8 Hz)** | P value | p < 0.001 | - |
| | t-value | 202.5 | - |
| | No. of electrode | 61 | - |
| **Alpha (8-13 Hz)** | P value | p < 0.001 | - |
| | t-value | 242.88 | - |
| | No. of electrode | 62 | - |
| **Low Beta (13-15 Hz)** | P value | p < 0.001 | - |
| | t-value | 211.05 | - |
| | No. of electrode | 61 | - |
| | P value | p < 0.001 | 0.0049 |

| | | | |
|---|---|---|---|
| **High Beta (23-32 Hz)** | t-value | 187.38 | 91.24 |
| | No. of electrode | 56 | 33 |
| **Low Gamma (32-50 Hz)** | P value | p < 0.001 | p < 0.001 |
| | t-value | 209.96 | 140.86 |
| | No. of electrode | 57 | 46 |
| **High Gamma (50-80 Hz)** | P value | p < 0.001 | p < 0.001 |
| | t-value | 217.24 | 198.4 |
| | No. of electrode | 59 | 58 |

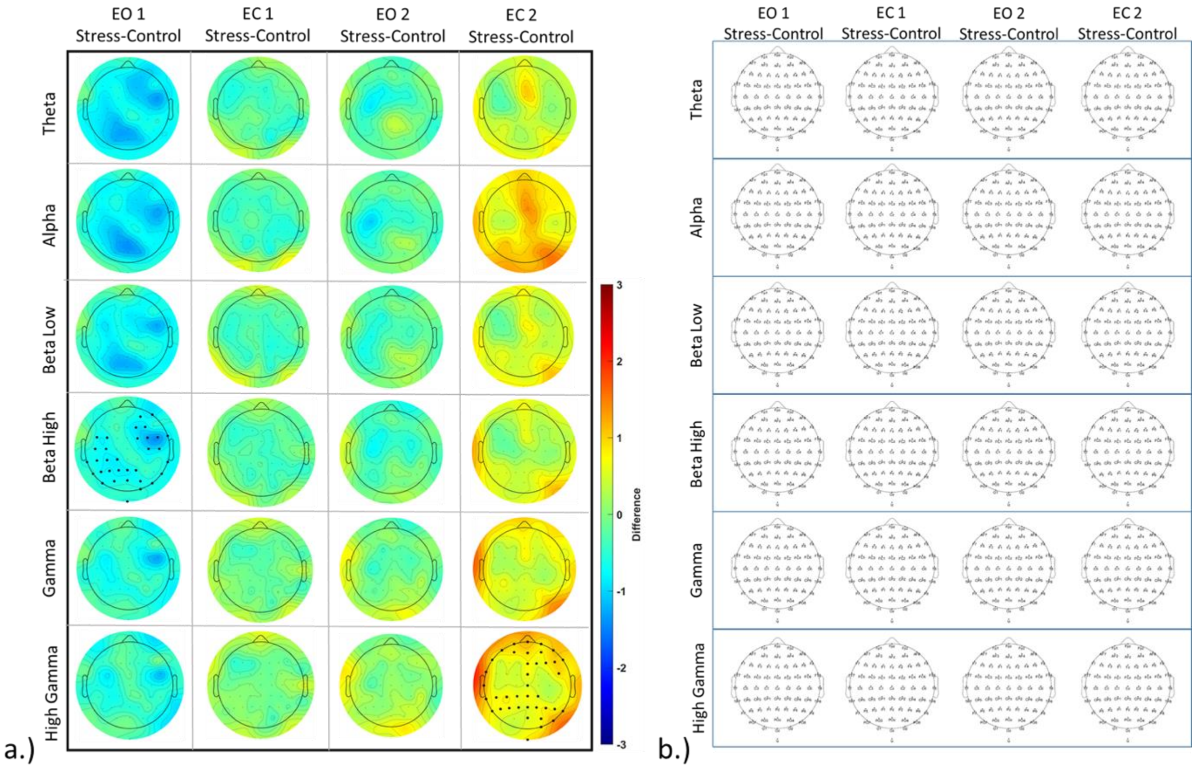


*Supplementary Figure S1 a.) EEG power changes during the resting state for both, control and stress clips. The topographical plots show power differences between control and stress conditions for each resting state in EO and EC conditions. The color bar indicates the difference value range from 3 to -3. Red shows a higher value and blue a lower value in the stress compared to the control intervention condition. Statistics were done as described in clip power statistics b.) EEG connectivity difference during resting state for both, control and stress conditions. No significant connectivity change was observed. Statistics were performed as described in clip connectivity statistics. EO denotes Eyes Open and EC Eyes Closed resting conditions.*
